# Strain-specific differences in *toxT* expression and virulence gene activation in *Vibrio cholerae*

**DOI:** 10.1101/2025.04.04.647171

**Authors:** Jayun Joo, Donghyun Lee, Seoyun Choi, Hyungoo Kim, Dong Wook Kim, Eun Jin Kim

## Abstract

The transcriptional activator ToxT is a key regulator of *Vibrio cholerae* virulence, controlling the expression of cholera toxin (CT) and the toxin co-regulated pilus (TCP). Insights into the regulation and function of *toxT* have largely come from studies on the classical biotype strain O395, which exhibits robust *toxT* expression. However, *toxT* expression is not fully consistent across *V. cholerae* strains. Here, we show that *toxT* transcription and protein production differ significantly among strains, particularly in El Tor biotype, which exhibit allele-dependent regulatory constraints. Using qRT-PCR and Western blotting, we demonstrate that alternative *toxT* alleles confer distinct transcriptional activator functions across El Tor biotype strains. Notably, IB5230, a pandemic El Tor strain, exhibits enhanced *toxT* and virulence gene expression even at 37°C, suggesting a potential adaptation for increased pathogenicity. Our results suggest that *toxT* autoregulatory feedback plays a more critical role in virulence gene activation than previously recognized. This study underscores previously unrecognized complexity in *toxT* regulation, emphasizing the need to reassess virulence control mechanisms in epidemic and emerging *V. cholerae* strains. Understanding these strain-specific differences is essential for refining models of cholera pathogenesis.

**Author Summary:** Cholera is a severe diarrheal disease caused by the bacterium *Vibrio cholerae*, which continues to pose a major threat to global health. The ability of *V. cholerae* to cause disease depends on the expression of two key virulence factors—cholera toxin (CT) and the toxin-coregulated pilus(TCP)— controlled by the transcriptional activator ToxT. While much of our understanding of ToxT regulation is based on classical biotype strains, most recent cholera outbreaks have been driven by El Tor biotype strains. In this study, we demonstrate that *toxT* expression and its ability to activate virulence genes vary markedly across *V. cholerae* strains, depending on both the specific *toxT* allele and the genetic background. Using isogenic strains engineered to express different *toxT* alleles, we show that some alleles are transcriptionally silent in certain El Tor strains while others are active, despite having similar functional capabilities when expressed. Notably, a hypervirulent strain linked to the Haitian cholera outbreak maintained high *toxT* expression even at human body temperature, hinting at possible adaptive mechanisms for enhanced pathogenicity. Our findings suggest that virulence regulation in *V. cholerae* may be more variable than previously recognized, highlighting strain- and allele-specific differences in *toxT* expression. Understanding these strain- and allele-specific differences in virulence gene regulation is crucial for improving models of cholera pathogenesis and could inform the development of more effective interventions and vaccines against this evolving pathogen.

## Introduction

*Vibrio cholerae* is a gram-negative bacterium and the etiological agent of cholera, a severe diarrheal disease that remains a major public health challenge, particularly in regions with inadequate sanitation and water supply systems [1, 2]. Based on the structure of its O-antigen, *V. cholerae* is classified into more than 200 distinct serogroups; however, strains belonging to the O1 and O139 serogroups are primarily responsible for epidemic cholera [1–3]. The clinically significant O1 serogroup is further subdivided into three serotypes: Ogawa, Inaba, and Hikojima [1].

Additionally, O1 serogroup strains are classified into two biotypes, classical and El Tor, based on distinct microbiological characteristics, including biochemical properties, phage susceptibility, and environmental resilience [2]. Historically, the classical biotype was responsible for the first six cholera pandemics recognized since the early 19^th^ century. In contrast, the ongoing seventh cholera pandemic, which began in 1961, represents a notable shift in disease epidemiology, as it has been predominantly attributed to El Tor biotype strains [3–5].

The ability of *V. cholerae* to cause disease is primarily driven by two key virulence factors: cholera toxin (CT) and toxin co-regulated pilus (TCP). CT, a potent enterotoxin secreted by strains within the epidemic serogroups O1 and O139, induces the profuse watery diarrhea characteristic of cholera, leading to severe dehydration and, if untreated, potentially fatal outcomes [6]. TCP plays a crucial role in the initial stages of infection by facilitating the colonization of the human small intestine, which is essential for the pathogen’s persistence and proliferation within the host.

*V. cholerae* relies on a complex regulatory network, the ToxR regulon, to ensure proper activation of CT and TCP in response to host environmental cues [7–11]. Central to this regulon is ToxT, an AraC-type transcriptional activator that directly regulates the expression of the virulence genes *ctxAB* and *tcpA*, which encode CT and the major structural component of TCP, TcpA, respectively [12].

ToxT functions by binding to specific DNA sequences within the promoter regions of these virulence genes, initiating their transcription. Experimental studies, including electrophoretic mobility shift assays (EMSA), reporter gene assays, and three-dimensional structural analyses, have demonstrated that ToxT directly activates the transcription of these target genes, leading to the subsequent production of CT and TCP [13–16].

Despite extensive research on virulence gene regulation, significant variability exists in virulence gene expression among different *V. cholerae* strains under laboratory conditions [17]. Classical biotype strains typically express virulence genes when cultured in aerated LB medium at pH 6.5 and 30°C (agglutinating conditions) [18], whereas El Tor biotype strains require the specialized AKI culture method, a two-phase process involving static incubation in AKI medium (composed of 1.5% Bacto peptone, 0.4% yeast extract, 0.5% NaCl, and 0.3% NaHCO_3_) at 37°C, followed by vigorous shaking [19]. However, some strains from both biotypes fail to express virulence genes even under their respective inducing conditions, suggesting the involvement of additional regulatory mechanisms [20, 21].

In most *V. cholerae* strains, including both classical and El Tor biotypes, the ToxT protein shares the same amino acid sequence and carries the *toxT*-SY allele, with Ser at position 65 and Tyr at position 139 [12]. However, recent studies have identified variant *toxT* alleles with distinct amino acid substitutions [20, 22]. The *toxT*-AY allele, found in the classical biotype strain 569B, is characterized by Ala at position 65 and Tyr at position 139. Meanwhile, the *toxT*-SF allele, present in the El Tor biotype strain MG116025, contains Ser at position 65 and Phe at position 139 [23]. Additionally, an artificial allele, *toxT*-AF, has been generated by combining the *toxT*-AY and *toxT*-SF alleles, resulting in Ala at position 65 and Phe at position 139.

Since the *toxT*-SF allele has been shown to facilitate virulence gene expression in several *V. cholerae* strains, previous studies have investigated the effects of replacing the authentic *toxT* allele with alternative alleles in a number of *V. cholerae* strains [20, 22, 23]. When introduced into various *V. cholerae* strains, these *toxT* alleles exhibited distinct effects on CT and TCP expression under laboratory culture conditions [20, 23]. In the classical biotype strain O395, all four alleles activated transcription of virulence genes [20]. However, in the El Tor biotype strain N16961, *toxT*-SY and *toxT*-SF failed to induce virulence gene expression under any tested condition, whereas *toxT*-AY and *toxT*-AF stimulated expression in aerated culture in LB medium [20, 24]. Notably, in another *V. cholerae* strain, IB5230, the strain responsible for the 2010 Haitian cholera outbreak, the regulatory pattern was entirely reversed: *toxT*-SY and *toxT*-SF promoted virulence gene expression, while *toxT*-AY and *toxT*-AF did not [22, 24, 25].

The variation in virulence gene expression across different strains and *toxT* alleles underscore the complexity of virulence gene regulation. This suggests that additional genetic or regulatory factors may influence virulence gene activation, highlighting previously unrecognized differences in genetic composition, regulatory networks, or environmental adaptability that impact pathogenic potential both *in vitro* and *in vivo*.

This study aimed to analyze whether differences in virulence gene expression among *V. cholerae* strains depend on the expression of *toxT* alleles. Specifically, we sought to assess whether these *toxT* alleles fail to be expressed or, alternatively, are expressed but unable to effectively activate downstream virulence genes in *V. cholerae* strain and *toxT* allele combinations that do not support virulence gene expression. To address this, we conducted a detailed analysis of *toxT*, *ctxAB*, and *tcpA* expression levels using qRT-PCR, further validated by Western blotting across *V. cholerae* strains.

Significant differences in *toxT* expression—and consequently, in the activation of CT and TCP production—were observed depending on the *toxT* allele and strain. Our results suggest that the regulation of virulence gene expression in *V. cholerae* is more complex and strain-dependent than previously understood.

## Results

### Functional characterization of *toxT* alleles

All four native *toxT* alleles exhibited transcriptional activator functions in the classical biotype strain O395 [20]. However, only specific *toxT* alleles (e.g., *toxT*-AY and *toxT*-AF in N16961, and *toxT*-SY and *toxT*-SF in IB5230) retained this function in other strains [20]. To confirm that all four *toxT* alleles maintain their transcriptional activator functions in El Tor biotype strains, we analyzed virulence gene expression by exogenously expressing *toxT* alleles in a *toxT*-deleted N16961 strain. An isogenic derivative of the N16961 strain, DHL010 (N16961-Δ*toxT*) was individually transformed with recombinant pBAD plasmids, each carrying a different His-tagged *toxT* allele [20]. The expression of His-tagged ToxT and TcpA was assessed in the presence of 0.2% arabinose or 0.5% glucose. The addition of glucose effectively suppressed basal expression of His-tagged ToxT. Since the O395 and IB5230 strains lose viability in the presence of 0.5% glucose due to impaired neutral fermentation, this experiment was performed exclusively in N16961 [20]. All four *toxT* alleles were successfully induced by the P_BAD_ promoter, leading to the expression of both His-tagged ToxT and TcpA in the presence of 0.2% arabinose. In contrast, neither ToxT nor TcpA was detected in the presence of 0.5% glucose (Fig. S1).

These results indicate that the *toxT* alleles exhibit no significant functional differences once expressed. This observation aligns with previous studies showing that all four *toxT* alleles in the O395 strain can activate virulence gene expression [20, 25].

### Functional equivalence of His-tagged ToxT compared to native ToxT

The introduction of a His-tag to *toxT* alters the amino acid sequence of both ToxT and the adjacent TcpJ protein, requiring an evaluation of the functionality of His-tagged ToxT (ToxT-His) relative to native ToxT (Fig. S2). To assess the potential impacts of these modifications, two complementary approaches were employed. First, the growth dynamics of isogenic derivatives expressing ToxT-His were compared to those of the parental strain expressing authentic ToxT under identical conditions. Second, the assembly and functionality of the TCP pilus, regulated by ToxT, were evaluated using CTX phage transduction efficiency. Together, these analyses provide a comprehensive assessment of the functional equivalence of ToxT-His and native ToxT, enabling an evaluation of the effects of His-tagging on ToxT activity and its downstream regulatory roles.

### Comparison of growth curves between isogenic derivatives harboring native *toxT* and His-tagged *toxT*

To evaluate whether the integration of a His-tag at the C-terminus of the chromosomal *toxT* gene affects growth in *V. cholerae* strains, growth curves were compared across four isogenic derivatives of each strain (Fig. S3). These included the parental strain containing the authentic *toxT* allele (*toxT*-SY), its derivative harboring the His-tagged *toxT* (*toxT*-SY-His), an isogenic derivative with an alternative *toxT* allele (*toxT*-AF), and its counterpart with the His-tagged alternative *toxT* allele (*toxT*-AF-His).

Bacteria were cultured in LB medium at pH 6.5 and 30°C, and their growth was monitored. After a brief lag phase, exponential growth was observed for up to six hours of cultivation, with cell densities reaching approximately 5 × 10^9^ CFU/ml at the stationary phase. The growth curves of the isogenic derivatives of each strain were nearly identical, indicating that replacing the authentic *toxT* with either a His-tagged *toxT* or an alternative *toxT* (including His-tagged alternative *toxT*) did not significantly affect the growth rate (Fig. S3).

### Production of functional TCP by alternative *toxT* alleles

The functional assembly of the TCP by TcpA, expressed from His-tagged ToxT, was evaluated by measuring CTX phage transduction efficiency [4, 23]. Transduction experiments were conducted using the CTX-1 phage (in which *ctxAB* was replaced by a kanamycin-resistance cassette), produced from pCTX-1 (the replicative form of the CTX phage), and four isogenic derivatives of the *V. cholerae* strain O395: O395 (*toxT*-SY), O395-H (His-tagged *toxT*-SY), EJK009 (*toxT*-AF), and EJK009-H (His-tagged *toxT*-AF), which were prepared under agglutinating conditions [18, 23, 26]. Transduction efficiency was determined as the ratio of transductants to the total number of recipient cells.

The observed transduction efficiencies were approximately 18.3% for O395, 15.9% for O395-H, 20.3% for EJK009, and 20% for EJK009-H (Table S1). However, when EJK009-H recipient cells were prepared in LB medium at 37°C—a condition known to repress virulence gene expression [18]—transduction efficiency dropped by more than 10³-fold. These results indicate that functional TCP was assembled, with no significant differences in transduction efficiency between isogenic derivatives expressing His-tagged ToxT and those expressing native ToxT. Furthermore, these results confirmed that the His-tag did not affect TcpA production, TCP assembly, or virulence gene regulation, indicating that His-tagged ToxT retained its ability to regulate virulence gene expression with minimal interference [23].

### Expression of native ToxT and His-tagged ToxT in the *V. cholerae* O395 strain

All four native *toxT* alleles (*toxT*-SY, *toxT*-SF, *toxT*-AY, and *toxT*-AF) have been shown to activate virulence gene expression under agglutinating conditions in the classical biotype strain O395 [20, 24]. To assess the functionality of native and His-tagged *toxT* variants, we compared the expression of *toxT* at the mRNA level by RT-PCR and at the protein level using Western blotting (anti-His-tag) in isogenic derivatives expressing the *toxT*-AF allele, with (EJK009-H) or without (EJK009) a His-tag, across different growth phases (Fig. 1). These isogenic derivatives were cultured under two conditions: agglutinating conditions (30°C in LB medium, pH adjusted to 6.5) and virulence-repressing conditions (LB medium at 37°C, pH adjusted to 8.5).

**Figure 1.**
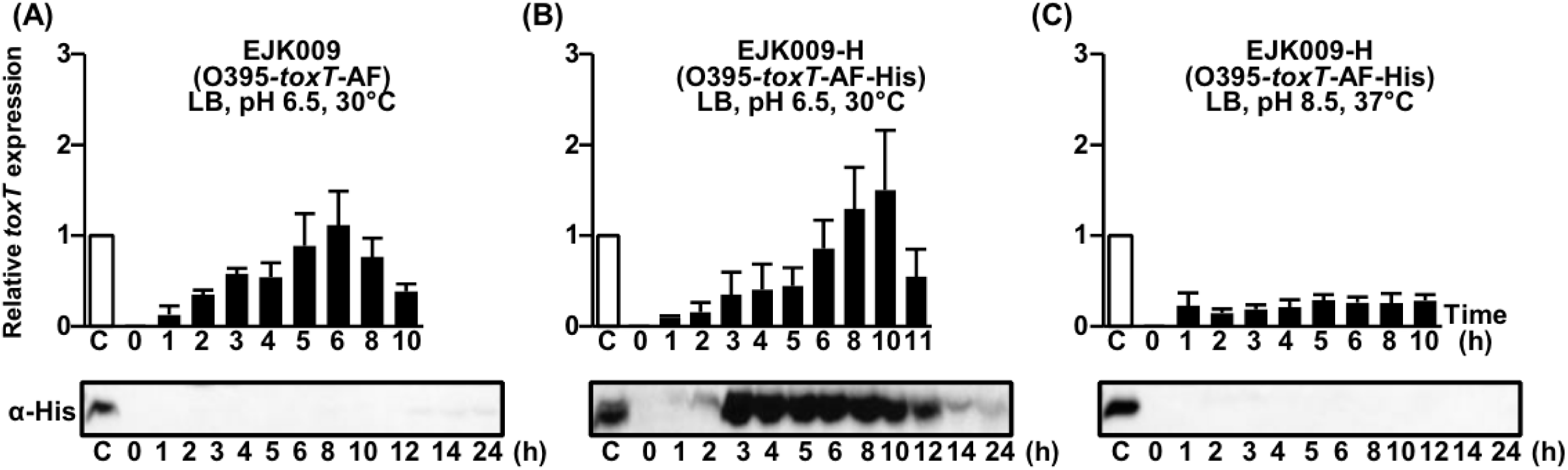
Comparison of native *toxT* and His-tagged *toxT* expression in EJK009 (O395-*toxT*-AF) and EJK009-H (O395-*toxT*-AF-His). qRT-PCR analysis and Western blot images of the isogenic derivatives probed with anti-His-tag antibodies. (A) Native *toxT* mRNA in EJK009 (O395-*toxT*-AF) cultured in LB medium (pH 6.5) at 30°C, (B) His-tagged *toxT* mRNA in EJK009-H (O395-*toxT*-AF-His) cultured in LB medium (pH 6.5) at 30°C, and (C) His-tagged *toxT* mRNA in EJK009-H cultured in LB medium (pH 8.5) at 37°C. Expression values were normalized to the housekeeping gene *gyrA*. The relative expression levels of *toxT* (black bar) at each time point are presented, with the *toxT* expression level in the 4-hour culture of O395-H (O395-*toxT-*SY-His, white bar) set to 1 (shown as lane C).

Since ToxT and TcpA of the O395 strain carrying the authentic *toxT* allele (*toxT*-SY) become detectable by Western blot starting at 4 hours after culture initiation under agglutinating conditions (described below), the mRNA levels of *toxT*, *tcpA*, and *ctxAB* of O395-H (His-tagged *toxT*-SY) at this time point were set as the reference (1, or 100%). The expression levels of each of the virulence gene were then compared relative to this reference across all strains based on their respective *toxT* alleles. Under agglutinating culture conditions, both EJK009 (*toxT*-AF) and EJK009-H (His-tagged *toxT*-AF) exhibited similar virulence gene expression patterns (Figs. 1 and 2). Detectable *toxT* mRNA was observed as early as 1–2 hours post-cultivation, peaking between 6 and 10 hours. Stable *toxT* mRNA levels persisted up to 10 hours, indicating consistent expression during this period. After 8-10 hours, *toxT* mRNA levels began to decline (Figs. 1A and 1B). ToxT protein was detectable by Western blot analysis as early as 2 hours post-cultivation. Under virulence-inducing conditions, ToxT protein levels peaked between 3 and 4 hours, gradually decreasing thereafter, with nearly undetectable levels after 14 hours (Fig. 1B). Under virulence-repressing conditions, only minimal levels of *toxT* mRNA were detected, and no detectable ToxT protein was observed (Fig. 1C).

**Figure 2.**
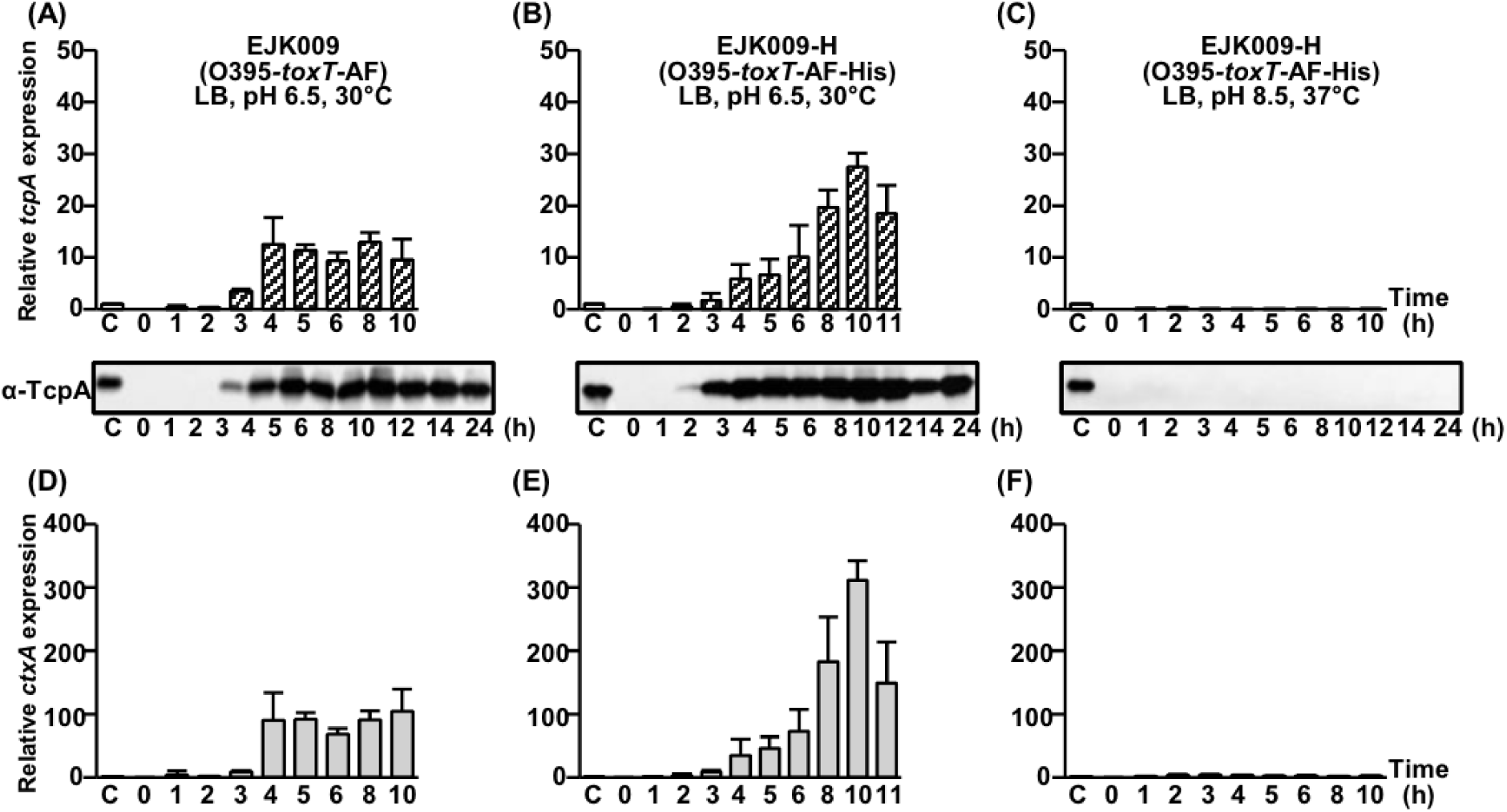
Comparison of *tcpA* and *ctxAB* expression in EJK009 and EJK009-H. qRT-PCR analysis of *tcpA* mRNA and Western blot detection of TcpA using anti-TcpA antibodies: (A) EJK009 cultured in LB medium (pH 6.5) at 30°C, (B) EJK009-H cultured in LB medium (pH 6.5) at 30°C, and (C) EJK009-H cultured in LB medium (pH 8.5) at 37°C. qRT-PCR analysis of *ctxAB* mRNA: (D) EJK009 and (E) EJK009-H cultured in LB medium (pH 6.5) at 30°C. (F) EJK009-H cultured in LB medium (pH 8.5) at 37°C. Expression values were normalized to the housekeeping gene *gyrA*. Lane C: O395-H (white bar), cultured in LB (pH 6.5) at 30°C for 4 hours. The relative expression levels of *tcpA* (diagonal bar) and *ctxAB* (gray bar) at each time point are presented, with the *tcpA* and *ctxAB* expression level in the 4-hour culture of O395-H set to 1.

These results indicate that the addition of a His-tag to *toxT* does not significantly alter *toxT* expression levels in *V. cholerae* strains under the conditions tested in this study.

### Virulence gene expression by native *toxT*-AF and His-tagged *toxT*-AF in the O395 strain

Having confirmed that the expression of His-tagged *toxT*-AF was comparable to that of native *toxT*-AF, we next investigated whether both native ToxT-AF and His-tagged ToxT-AF similarly activated the expression of downstream target genes, *tcpA* and *ctxAB*. The expression of these target genes was observed to follow a slight delay relative to *toxT* expression.

Under agglutinating culture conditions, *tcpA* and *ctxAB* mRNA became detectable approximately 3 hours after cultivation, with TcpA protein also becoming detectable at this time. TcpA protein levels were maintained up to 24 hours (Figs. 2A and 2B). However, under virulence-repressing conditions, no expression of *tcpA* or *ctxAB* was detected (Figs. 2C and 2F). These results demonstrate that the addition of the His-tag does not significantly affect the function of ToxT as a transcriptional activator of virulence genes.

### Virulence gene expression by alternative *toxT* alleles in the O395 strain

Given that the expression of His-tagged *toxT*-AF and consequently *tcpA* and *ctxAB*, was comparable to that of native *toxT*-AF in the O395 strain, we hypothesized that the virulence gene expression pattern in isogenic derivatives carrying the His-tagged *toxT* allele could serve as a surrogate for the corresponding native *toxT* allele. Hereafter, we investigated the expression of virulence genes in these isogenic derivatives harboring His-tagged *toxT* alleles.

#### toxT-SY

The O395 strain harbors the *toxT*-SY allele. Under agglutinating conditions, *toxT* mRNA in O395-H (His-tagged *toxT*-SY) was detectable within two hours of cultivation, reaching its highest levels at 5 hours (Fig. S4A). His-tagged ToxT protein was first observed at 3 hours, with protein levels peaking between 3 and 4 hours (Fig. S4A). Both *toxT* mRNA and ToxT protein levels began to decline after reaching their peaks, with *toxT* mRNA decreasing by 8 hours and ToxT protein levels dropping by 12 hours. Under virulence-repressing conditions, *toxT* mRNA levels were minimal, and ToxT protein production was negligible (Figs. S4B).

The expression of *tcpA* and *ctxAB* mRNA was first detected between 2 and 4 hours after cultivation, with peak expression occurring at approximately 5–6 hours, followed by a gradual decrease (Figs. S4C and S4E). TcpA protein levels became detectable starting at 4 hours, peaked between 7 and 8 hours, and remained detectable up to 24 hours post-cultivation (Fig. S4C). No expression of *tcpA* and *ctxAB* was detected under virulence-repressing conditions (Figs. S4D and S4F).

#### toxT-SF

The expression of His-tagged *toxT*-SF in YJB001-H under agglutinating conditions was first detectable within one hour of culture, peaking at 2 hours before gradually decreasing (Fig. S5A). His-tagged ToxT was detectable starting at one hour post-cultivation, with peak expression occurring between 2 and 3 hours (Fig. S5A). In contrast, under virulence-repressing conditions, *toxT* mRNA levels were minimal, and His-tagged ToxT was undetectable (Fig. S5B).

The mRNAs of *tcpA* and *ctxAB* were first detected at 3 hours post-cultivation, reaching peak levels at 8 hours, followed by a decline (Figs. S5C and S5E). TcpA protein levels were first detectable at 4 hours, with peak levels observed between 10 and 12 hours. TcpA protein levels remained substantial for up to 24 hours (Fig. S5C). Under virulence-repressing conditions, no expression of *tcpA* or *ctxAB* was observed (Figs. S5D and S5F).

#### toxT-AY

The expression of His-tagged *toxT*-AY mRNA in EJK008-H (His-tagged *toxT*-AY) was first detected at 2 hours post-cultivation under agglutinating conditions, with mRNA levels reaching a peak at 5 hours before progressively declining (Fig. S6A). His-tagged ToxT protein was similarly detected starting at 2 hours (Fig. S6A). Under virulence-repressing conditions, both *toxT*-AY mRNA and His-tagged ToxT protein were present at negligible levels (Fig. S6B).

Under agglutinating conditions, *tcpA* and *ctxAB* mRNA became detectable at 2 hours post-cultivation, peaking at 4–5 hours before gradually declining (Figs. S6C and S6E). TcpA protein was first detected at 3 hours, with substantial levels remaining detectable for up to 24 hours post-cultivation (Fig. S6C). Under virulence-repressing conditions, the expression of *tcpA* and *ctxAB* was negligible (Figs. S6D and S6F).

### Virulence gene expression in El Tor biotype strains

We investigated two *V. cholerae* El Tor biotype strains: the Wave 1 prototype strain N16961 and the Wave 3 atypical El Tor strain IB5230 [3, 27]. Although El Tor biotype strains are generally known to express virulence genes under AKI conditions, our laboratory observations indicated that while IB5230 did express these genes under such conditions, N16961 did not [20, 22, 25]. However, when alternative *toxT* alleles (*toxT*-AY and *toxT*-AF) were introduced in N16961, virulence gene expression was observed in aerated cultures using LB medium, even without pH adjustment to 6.5 [20, 21]. We monitored virulence gene expression of El Tor biotype strains grown in LB medium without pH adjustment.

#### N16961

The *V. cholerae* N16961 strain carries the authentic *toxT*-SY allele but does not exhibit virulence gene expression, even under AKI conditions [20, 25]. An isogenic derivative in which *toxT*-SY was replaced with *toxT*-SF also failed to induce virulence gene expression [20]. However, previous studies have shown that isogenic derivatives in which *toxT*-SY was replaced with *toxT*-AY or *toxT*-AF exhibited virulence gene expression at 30°C [20].

In this study, we investigated virulence gene expression in isogenic derivatives of N16961 carrying four different His-tagged *toxT* alleles. The isogenic derivatives carrying His-tagged *toxT*-AF (DHL009-H) or His-tagged *toxT*-AY (DHL008-H) exhibited virulence gene expression during the early exponential phase at 30°C (Figs. 3A, 4A, S7A, C, and E). Once mRNA transcription was initiated, ToxT protein was produced immediately and remained detectable for approximately 12 hours (Fig. 3A). *tcpA* and *ctxAB* expression followed a similar pattern (Fig 4A and 4E).

**Figure 3.**
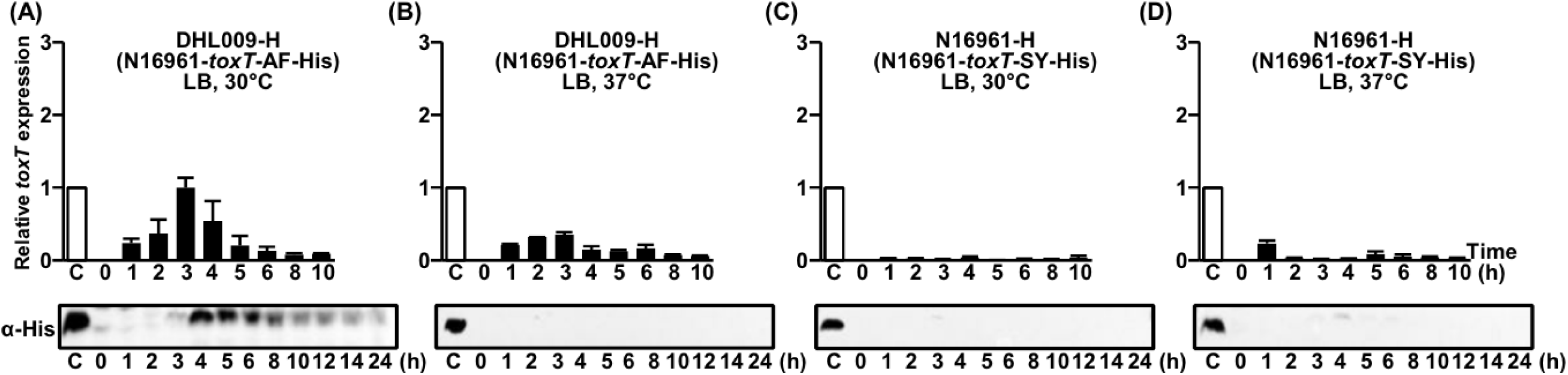
Analysis of *toxT* expression in DHL009-H (N16961-*toxT*-AF-His) and N16961-H (N16961-*toxT*-SY-His). qRT-PCR analysis of His-tagged *toxT* mRNA and Western blot detection of ToxT using anti-His-tag antibodies in (A) DHL009-H (N16961-*toxT*-AF-His) cultured in LB medium at 30°C, (B) DHL009-H cultured in LB at 37°C, (C) N16961-H (N16961-*toxT*-SY-His) cultured in LB at 30°C, and (D) N16961-H cultured in LB at 37°C. Expression values were normalized to the housekeeping gene *gyrA*. Lane C: O395-H (white bar), cultured in LB (pH 6.5) at 30°C for 4 hours. The relative expression levels of *toxT* (black bar) at each time point are presented, with the *toxT* expression level in the 4-hour culture of O395-H set to 1.

**Figure 4.**
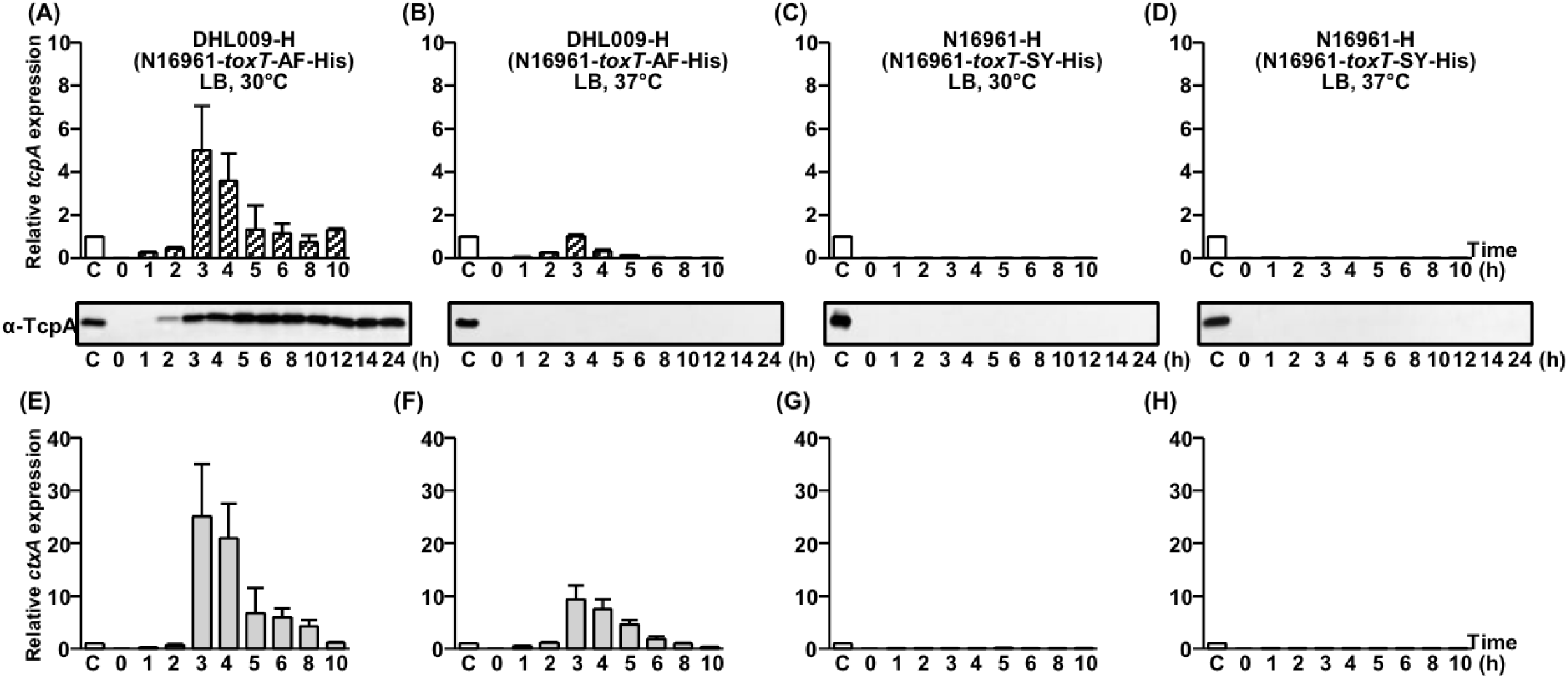
Comparison of *tcpA* and *ctxAB* expression in DHL009-H and N16961-H. qRT-PCR analysis of *tcpA* mRNA and Western blot detection of TcpA using anti-TcpA antibodies in (A) DHL009-H cultured in LB medium at 30°C, (B) DHL009-H cultured in LB at 37°C, (C) N16961-H cultured in LB at 30°C, and (D) N16961-H cultured in LB at 37°C. qRT-PCR analysis of *ctxAB* mRNA in (E) DHL009-H cultured in LB at 30°C, (F) DHL009-H cultured in 37°C, (G) N16961-H cultured in LB at 30°C, and (H) N16961-H cultured in LB at 37°C. Expression values were normalized to the housekeeping gene *gyrA*. The relative expression levels of *tcpA* (diagonal bar) and *ctxAB* (gray bar) at each time point are presented, with the expression levels of *tcpA* and *ctxAB* in the 4-hour culture of O395-H in LB medium (pH 6.5) at 30°C set to 1 (lane C, white bar).

In contrast, no virulence gene expression was observed in isogenic derivatives carrying His-tagged *toxT*-SY (N16961-H) or His-tagged *toxT*-SF (YJB003-H) alleles (Figs. 3C, S8A, C, and E). Additionally, none of the *toxT* alleles supported virulence gene expression at 37°C (Figs. 3B, 3D, S8B, D, and F). Overall, the *tcpA* and *ctxAB* expression patterns observed with His-tagged *toxT* alleles were consistent with those observed with native *toxT* alleles [20, 25].

#### IB5230

The *V. cholerae* IB5230 strain also carries the authentic *toxT*-SY allele. Unlike N16961, IB5230 exhibited virulence gene expression when harboring either the *toxT*-SY or *toxT*-SF allele. However, replacing the authentic *toxT*-SY allele with the *toxT*-AY or *toxT*-AF allele abolished virulence gene expression [20, 25].

When the His-tagged *toxT*-SF allele was introduced into IB5230 (YJB020-H) and cultured at 30°C, *toxT* mRNA became detectable starting 1 hour after inoculation, reaching peak levels at approximately 4–5 hours (Fig. 5A). The ToxT protein also appeared around 1–2 hours after inoculation, with substantial levels maintained for approximately 10 hours. The expression of *tcpA* and *ctxAB* followed immediately after *toxT* expression (Fig. 6A and 6E).

**Figure 5.**
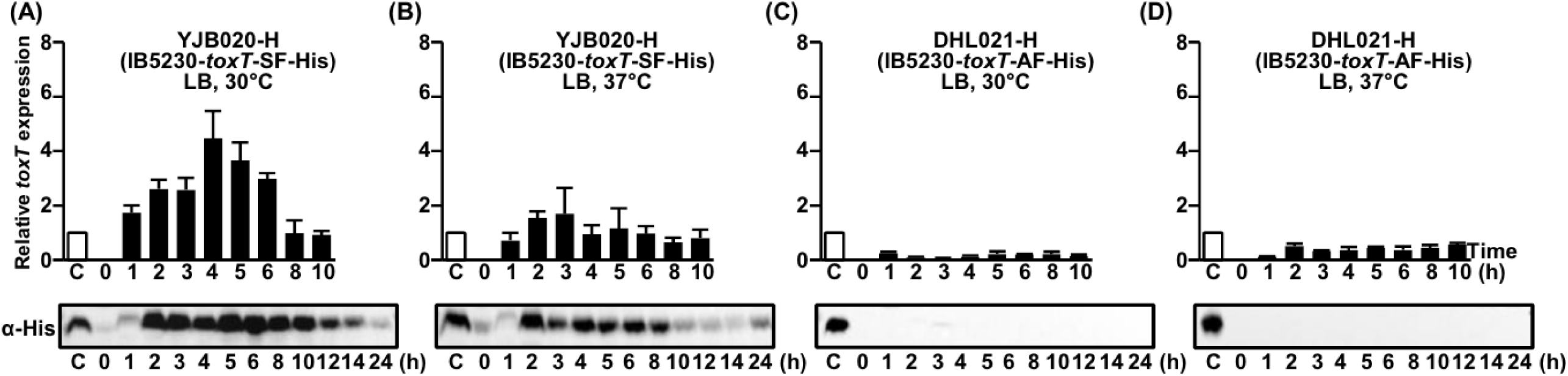
Comparison of *toxT* expression in YJB020-H (IB5230-*toxT*-SF-His) and DHL021-H (IB5230-*toxT*-AF-His). qRT-PCR analysis of His-tagged *toxT* mRNA and Western blot detection of ToxT using anti-His-tag antibodies in (A) YJB020-H (IB5230-*toxT*-SF-His) cultured in LB medium at 30°C, (B) YJB020-H cultured in LB at 37°C, (C) DHL021-H (IB5230-*toxT*-AF-His) cultured in LB at 30°C, and (D) DHL021-H cultured in LB at 37°C. Expression values were normalized to the housekeeping gene *gyrA*. The relative expression levels of *toxT* (black bar) at each time point are presented, with the *toxT* expression level in the 4-hour culture of O395-H in LB (pH 6.5) at 30°C set to 1 (Lane C, white bar).

**Figure 6.**
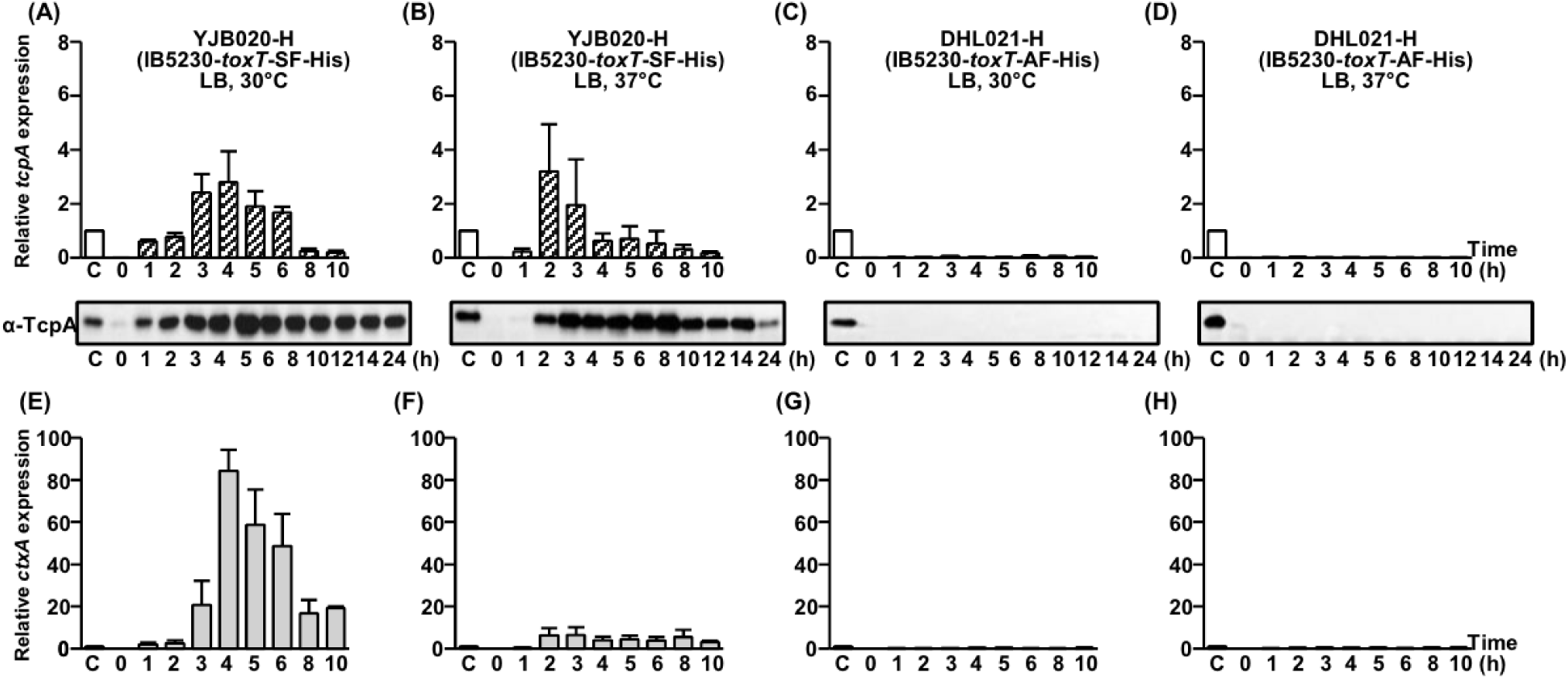
Comparison of *tcpA* and *ctxAB* expression in YJB020-H and DHL021-H. qRT-PCR analysis of *tcpA* mRNA and Western blot detection of TcpA using anti-TcpA antibodies in (A) YJB020-H (IB5230-*toxT*-SF-His) cultured in LB medium at 30°C, (B) YJB020-H cultured in LB at 37°C, (C) DHL021-H (IB5230-*toxT*-AF-His) cultured in LB at 30°C, and (D) DHL021-H cultured in LB at 37°C. qRT-PCR analysis of *ctxAB* mRNA in (E) YJB020-H cultured in LB at 30°C and (F) YJB020-H cultured in LB at 37, (G) DHL021-H cultured in LB at 30°C and (H) DHL021-H cultured in LB at 37°C. Expression values were normalized to the housekeeping gene *gyrA*. The relative expression levels of *tcpA* (diagonal bar) and *ctxAB* (gray bar) at each time point are presented, with the expression levels of *tcpA* and *ctxAB* in the 4-hour culture of O395-H in LB (pH 6.5) at 30°C set to 1 (lane C, white bar).

Unlike O395 or N16961, in which *toxT* and virulence gene expression were not observed at 37°C, YJB020-H exhibited *toxT* expression even at this temperature. At 37°C, *toxT* mRNA was detectable as early as 1 hour after inoculation, reaching its peak at approximately 3 hours. The ToxT protein appeared between 1 and 2 hours but declined more rapidly compared to the 30°C culture condition (Fig. 5B).

The expression of *tcpA* and *ctxAB* appeared to be differentially regulated. While *tcpA* mRNA and TcpA protein were detected (Fig. 6B), *ctxAB* expression was not observed (Fig. 6F). These results were consistent with those obtained from YJB020, in which the native *toxT*-SF allele without the His-tag was introduced [20, 25].

The isogenic derivative of IB5230 harboring the His-tagged *toxT*-AF allele (DHL021-H) did not exhibit *toxT*, *tcpA*, or *ctxAB* expression at either 30°C or 37°C (Fig. 5C, 5D, and Fig. 6C, D).

In the isogenic derivative of IB5230 harboring the His-tagged *toxT*-SY allele (IB5230-H), a temperature-dependent shift in *toxT* and virulence gene expression was observed. At 30°C, *toxT*, *tcpA*, and *ctxAB* were not expressed (Fig. S9A, C, and E). However, when cultured at 37°C, *toxT* and *ctxAB* expression were detected. Interestingly, *tcpA* expression remained absent under these conditions (Fig. S9B, D, and F).

In DHL020-H, which harbors the His-tagged *toxT*-AY allele, the expression of *toxT*, *tcpA*, and *ctxAB* was negligible at both 30°C and 37°C (Fig. S10A–F).

The expression patterns of *tcpA* and *ctxAB* in the isogenic derivatives of IB5230 harboring His-tagged *toxT* alleles—where virulence gene expression was observed at 37°C or where *tcpA* and *ctxAB* were differentially expressed—closely matched those observed when native *toxT* alleles without a His-tag were introduced [20]. These results suggest a potential unique virulence gene expression pattern in this strain.

### Inhibition of expression of ToxT by unsaturated fatty acids

The activity of ToxT is known to be regulated by fatty acids present in bile [28]. While saturated fatty acids (SFAs), such as stearic acid, have little to no effect on ToxT function as a transcriptional activator of virulence genes, unsaturated fatty acids (UFAs), including linoleic acid, have been reported to suppress ToxT activity [25, 29, 30].

Previous studies have demonstrated that UFAs, particularly linoleic acid, can enter *V. cholerae* cells and directly reduce ToxT’s ability to bind to DNA at virulence promoters [12, 14, 29]. This reduction in target DNA binding leads to decreased expression of virulence genes. These findings suggested that UFAs inhibit ToxT monomers from binding DNA—a mechanism distinct from that of virstatin, a synthetic ToxT inhibitor, which prevents ToxT dimerization [12, 29, 31].

The present study aimed to further investigate this regulatory mechanism by comparing the ToxT and TcpA production in the presence of different fatty acids. Specifically, we sought to determine whether UFAs inhibit ToxT synthesis or act after ToxT has already been synthesized.

The results indicate that, when UFAs are present in the culture medium, ToxT production is significantly reduced, whereas saturated fatty acids do not cause noticeable changes (Fig. 7A). Consequently, TcpA expression is almost completely abolished in the presence of UFAs (Fig. 7B). These findings suggest that UFAs inhibit ToxT at the production stage rather than solely at the functional level. Furthermore, these results suggest that UFAs may bind to the initially synthesized ToxT through the ToxR regulon in response to external stimuli, thereby disrupting its autoregulatory positive feedback loop and leading to reduced ToxT expression (discussed below).

**Figure 7.**
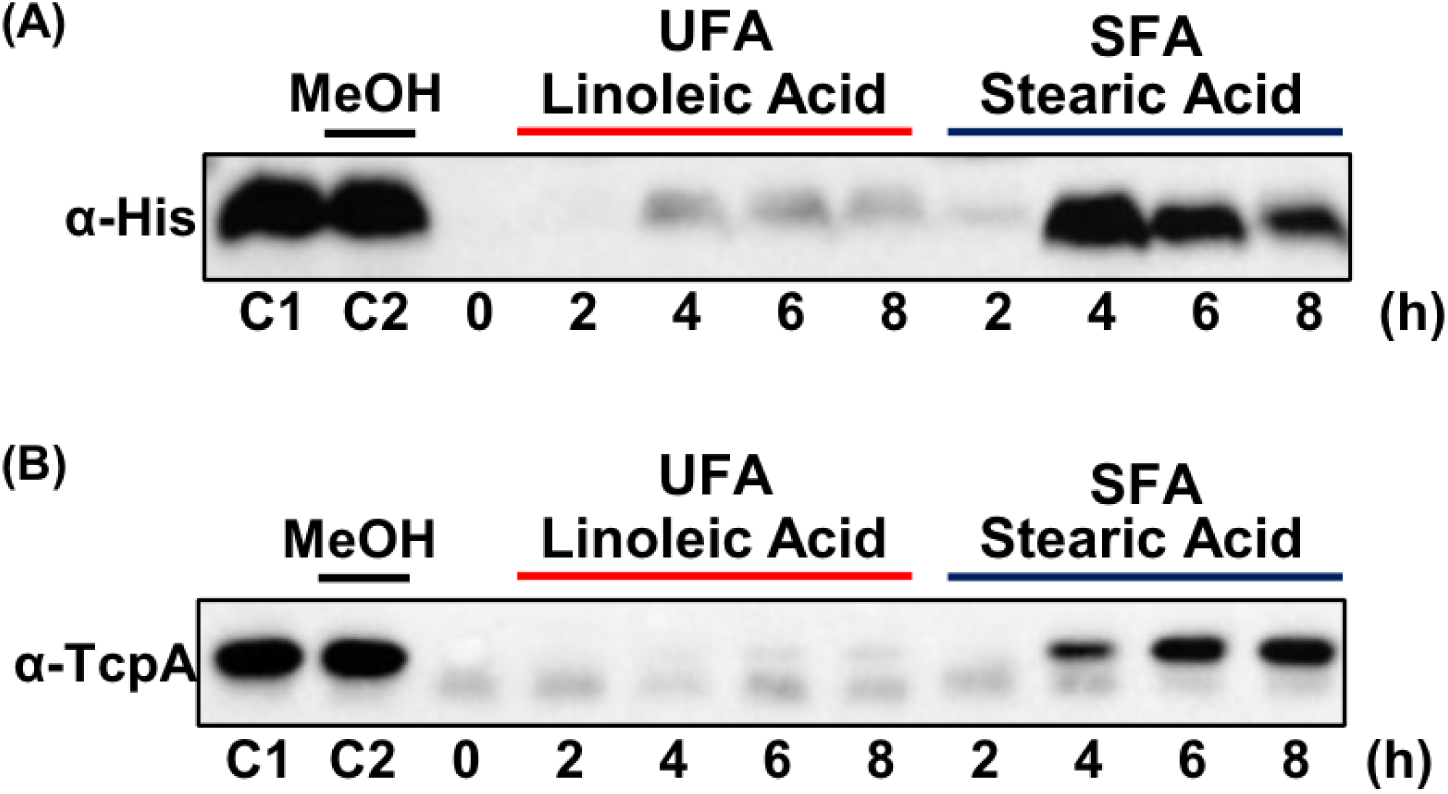
Impact of fatty acids on the expression of His-tagged ToxT and TcpA. Western blot analysis of His-tagged ToxT in O395-H (*toxT-*SY-His) using (A) anti-His-tag antibodies and (B) anti-TcpA antibodies. The bacteria were cultured in LB medium (pH 6.5) at 30°C for up to 8 hours in the presence of linoleic acid (unsaturated fatty acids, UFA) or stearic acid (SFA, saturated fatty acids). Lane C1: O395-H, cultured for 4 hours; Lane C2: O395-H, cultured for 4 hours in the presence of methanol (1/100 culture volume). The final concentration of the fatty acids was 0.02%.

## Discussion

The two primary virulence genes of *V. cholerae*, *ctxAB* and *tcpA*, are regulated by the ToxR regulon, with ToxT serving as the final transcriptional activator [32]. While *toxT* expression and its regulatory mechanisms have been extensively studied in the classical biotype strain O395, virulence gene expression in El Tor biotype strains remains less well understood. Although specific culture conditions have been developed to induce virulence gene expression and aid in understanding the pathogenicity of *V. cholerae*, these conditions cannot be universally applied to all *V. cholerae* strains [20–22].

Previously, we demonstrated that replacing the authentic *toxT* allele with alternative *toxT* alleles can alter virulence gene expression in *V. cholerae* strains [5, 20–22, 33]. Depending on the strain, certain *toxT* alleles can either induce or fail to induce virulence gene expression [20]. This study focused on the strain-dependent expression of *toxT* alleles in *V. cholerae* and their impact on virulence gene expression [34, 35]. To examine the expression of *toxT* alleles and their impacts on virulence gene expression, the native *toxT* allele in each isogenic derivative of *V. cholerae* was replaced with its corresponding His-tagged version.

In the classical biotype strain O395, all four His-tagged *toxT* alleles were expressed and activated virulence gene expression, although the level of activation varied. In contrast, the El Tor biotype strains exhibited His-tagged *toxT* allele-specific regulation. In N16961, His-tagged *toxT*-AY and *toxT*-AF were expressed and activated virulence gene transcription when the isogenic derivatives were cultured under simple shaking conditions in LB medium at 30°C. However, His-tagged *toxT*-SY and *toxT*-SF were not expressed, leading to the absence of virulence gene expression. The opposite pattern was observed in IB5230: His-tagged *toxT*-SY and *toxT*-SF were expressed and capable of activating virulence gene transcription, while His-tagged *toxT*-AY and *toxT*-AF were not expressed, resulting in a failure to induce virulence gene expression.

Compared to previous reports on virulence gene expression regulated by alternative *toxT* alleles in *V. cholerae* strains [20], the introduction of a His-tag to the chromosomal *toxT* did not alter the virulence gene expression pattern. *toxT* alleles capable of inducing virulence gene expression did so regardless of the presence of the His-tag, while *toxT* alleles that were unable to induce virulence gene expression remained inactive irrespective of the presence of the His-tag. Therefore, we inferred that the detection of His-tagged ToxT reflected the kinetics of native ToxT under laboratory culture conditions.

Regardless of the strain, when *toxT* expression occurred, *toxT* mRNA and ToxT protein were detectable from the early exponential phase and remained at substantial levels at least until the early stationary phase. This stability suggests that ToxT is persistent without significant fluctuations throughout the exponential and early stationary growth phases. ToxT levels may persist unless abrupt changes in the culture environment or the introduction of external factors specifically repress virulence gene expression. Furthermore, *tcpA* and *ctxAB* were immediately expressed following ToxT activation, leading to the production of functional TCP and CT.

In El Tor biotype strains, *toxT* alleles that failed to induce virulence gene expression were transcriptionally repressed, resulting in undetectable ToxT protein and, consequently, no expression of *ctxAB* and *tcpA*. These findings highlight the regulatory mechanisms governing ToxT expression in the absence of external repressive signals. Moreover, these results suggest that the expression of specific *toxT* alleles varies under identical culture conditions, and that the activation or repression of a particular *toxT* allele is differentially regulated depending on the *V. cholerae* strain.

Although the classical biotype strain O395 exhibited virulence gene activation with all *toxT* alleles, this trait is not shared by all classical biotype strains. Other classical strains, such as 569B, Cairo48, and Cairo50, displayed *toxT* allele-specific virulence gene expression, similar to that observed in El Tor biotype strains.

For example, in the 569B strain, the authentic *toxT*-AY allele supports virulence gene expression, as does the alternative *toxT*-AF allele. However, replacing these with *toxT*-SY or *toxT*-SF abolishes virulence gene activation [20, 22]. Similarly, the Cairo48 and Cairo50 strains, both used in oral cholera vaccine (OCV) development, naturally harbor the *toxT*-SY allele and fail to express virulence genes under agglutinating culture conditions. However, virulence gene expression could be restored when *toxT*-SY is replaced with alternative alleles—*toxT*-SF and *toxT*-AY in Cairo48, and *toxT*-AY in Cairo50 [20, 22]. These findings suggest that O395’s ability to express virulence genes independently of *toxT* allele type is a unique characteristic of this strain rather than a general feature of classical biotype strains.

All four *toxT* alleles were capable of activating virulence gene expression in the classical biotype strain O395, indicating that the function of ToxT itself does not differ significantly across alleles. This was further supported by our results that all four *toxT* alleles, when expressed exogenously under the P_BAD_ promoter in an isogenic derivative of the N16961 strain lacking the authentic *toxT* gene, successfully induced the expression of TcpA and CT. These results suggest that, while all *toxT* alleles can function as transcriptional activators in *V. cholerae* strains, their expression levels may serve as the primary determinant of virulence gene activation.

This raises a key question: why do some *toxT* alleles become transcriptionally active while others remain silent in a strain-dependent manner? One possible regulatory influence emerges from our study on the effects of unsaturated fatty acids (UFAs) on *toxT* expression. UFAs have been shown to inhibit ToxT function by preventing its binding to target DNA [12, 14]. Our results indicated that UFAs also suppress *toxT* expression itself. Since *toxT* expression in *V. cholerae* is regulated through a positive autoregulatory loop [36, 37], our results suggest a potential mechanism in which *toxT* is initially transcribed through the ToxR regulon but fails to establish its own positive feedback loop due to UFA interference. Consequently, ToxT accumulation remains insufficient for full transcriptional activation.

Occasionally, we have observed negligible or low levels of *toxT* mRNA and faint western blots (Fig. S4B, Fig. 7A), which may reflect primary or sub-threshold ToxT expression. Based on this, we hypothesize that a basal, undetectable level of ToxT may exist under our experimental conditions.

Once a certain threshold is reached through autoregulatory amplification, ToxT may become functionally active in virulence gene expression. It remains to be investigated whether this low-level expression of ToxT is an integral part of the ToxR regulon, representing primary ToxT production before the subsequent autoregulatory positive feedback loop.

Additionally, the factors that enable specific *toxT* alleles to engage in this autoregulatory loop while others remain inactive remain unclear. As proteolytic degradation of ToxT plays a critical role during the termination of virulence gene expression, it is possible that ToxT degradation also contributes significantly during the activation phase of virulence [38, 39]. To address this, future studies should incorporate more accurate quantitative approaches to distinguish between primary *toxT* expression and autoregulatory *toxT* expression.

Notably, IB5230, which was associated with the 2010 Haitian cholera outbreak, exhibited robust expression of *toxT* and virulence genes even at 37°C. This may align with the physiological conditions encountered in the human intestine, suggesting that strain-specific differences in *toxT* regulation could influence colonization efficiency and pathogenicity. Given that IB5230 and other wave 3 atypical El Tor strains have been globally prevalent as hypervirulent strains since the 2000s, further research is needed to determine whether *toxT* alleles or other regulatory mechanisms have evolved to enhance adaptation to human infection [5, 40, 41].

Although our findings were obtained under laboratory conditions and may not fully reflect the *in vivo* environment, they provide important insights into the regulatory mechanisms governing virulence gene expression in *V. cholerae*. Our results suggest that *toxT* allele- and strain-specific regulation is more complex and finely controlled than previously thought. Further studies are needed to determine whether the strain-dependent *toxT* activation patterns observed *in vitro* directly correlate with virulence gene expression and disease progression in the host. Elucidating the genetic and environmental factors that influence *toxT* activation will be essential for developing a more comprehensive understanding of virulence regulation during infection.

## Materials and Methods

### Bacterial strains

1. *V. cholerae* strains and their isogenic derivatives are listed in, Table S2.

### *toxT* Allele exchange

#### Allele exchange at the 139^th^ amino acid position

The replacement of the chromosomal authentic *toxT* with alternative *toxT* alleles into *V. cholerae* strains has been described previously [4, 20, 23]. Briefly, two 843 bp DNA fragments, encompassing the 50 nucleotides upstream of the translation start codon to nucleotide 793 of *toxT*, were amplified by PCR using the primers toxT-XbaIF: CCG GCC <u>TCT AGA</u> TAC GTG GAT GGC TCT CTG CG and toxT-SacIR: CCG GCC <u>GAG CTC</u> CAC TTG GTG CTA CAT TCA. These fragments were derived from the genomic DNA of *V. cholerae* strains MG116025 and N16961, respectively. The amplified DNA fragments were inserted into the suicide plasmid pCVD442 [42] to construct pCVD-toxT-SF and pCVD-toxT-SY, respectively. A single nucleotide polymorphism (SNP) at position 416 (A416 in N16961 and T416 in MG116025) is located centrally within these fragments. Using the allelic exchange method, the *toxT-SF* allele from the MG116025 strain was replaced with the *toxT-SY* allele from the N16961 strain, and conversely, the *toxT-SY* alleles in other strains were replaced with the *toxT-SF* allele from the MG116025 strain [4, 23]. Additionally, the 569B classical biotype strain contains another variant of *toxT*, referred to as *toxT*-AY. The pCVD-toxT-SF plasmid was conjugally transferred into 569B, resulting in the construction of a variant strain containing the *toxT-AF* allele through allelic exchange. Validation of the replaced *toxT* allele, following the excision of the pCVD442 suicide vector, was performed using Sanger sequencing.

#### Allele exchange at the 65^th^ amino acid position

A set of allele exchange plasmids was utilized to construct isogenic derivatives of *V. cholerae* strains harboring SNPs at the 65^th^ and 139^th^ amino acid positions of *toxT*. Four 1308 bp DNA fragments, spanning from the 709^th^ nucleotide of *tcpF* to the 793^rd^ nucleotide of *toxT*, were PCR-amplified using the primer pair TcpF-XbaIF: GGG <u>TCT AGA</u> GAA TTA AGT AAG CAC GGG TA and toxT-SacIR: CCG GCC <u>GAG CTC</u> CAC TTG GTG CTA CAT TCA. These fragments were obtained from the genomic DNA of the N16961, YJB003 (N16961-*toxT*-SF), 569B, and EJK007 (569B-*toxT*-AF) strains [21].

The amplified DNA fragments were subcloned into the suicide plasmid pCVD442 to construct the plasmids pCVD-toxT-Out-SY, pCVD-toxT-Out-SF, pCVD-toxT-Out-AY, and pCVD-toxT-Out-AF. These plasmids were conjugally transferred into *V. cholerae* strains to construct isogenic derivatives harboring alternative *toxT* alleles through the allelic exchange method. The excision of the suicide vector was screened on sucrose-containing agar plates, and the DNA sequences of the variant *toxT* alleles were confirmed by Sanger sequencing.

### Construction of a *toxT-*deleted (Δ*toxT*) isogenic variant of *V. cholerae* strains

A 635-bp DNA fragment, spanning from the 709^th^ nucleotide of *tcpF* to the 120^th^ nucleotide of *toxT*, was PCR-amplified using the primer pair TcpF-KpnIF: GGG <u>GGT ACC</u> GAA TTA AGT AAG CAC GGG TA and TcpF-BamHIR: CCC <u>GGA TCC</u> TGC AAT TCC ACT ATC TAT CCA [20]. This fragment was derived from the genomic DNA of *V. cholerae* strain N16961 and inserted into the KpnI and BamHI restriction sites of pUC18 to construct pUC-tcpF. Next, a 660-bp DNA fragment, spanning from the 616^th^ nucleotide of *toxT* to the 435^th^ nucleotide of *tcpJ*, was PCR-amplified using the primer pair TcpJ-BamHIF: GGG <u>GGA TCC</u> CTA GAG TCT CGA GGA GTA AAG and TcpJ-PstIR: CCC <u>CTG CAG</u> GCT GCA GAT AAA TAA ATG CC. This fragment was also derived from the genomic DNA of the N16961 strain and inserted into the BamHI and PstI sites of pUC-tcpF, resulting in the construction of pUC-tcpF-tcpJ. A primer pair TcpF-XbaIF: GGG <u>TCT AGA</u> GAA TTA AGT AAG CAC GGG TA and TcpH-SacIR: CCC <u>GAG CTC</u> GCT GCA GAT AAA TAA ATG CC was used to PCR-amplify a 1301-bp DNA fragment from pUC-tcpF-tcpJ, and this DNA fragment was inserted into the suicide plasmid pCVD442, generating pCVD-del-toxT. The pCVD-del-toxT plasmid was conjugally transferred into *V. cholerae* strains to construct *toxT*-deleted (Δ*toxT*) isogenic derivatives. Following the excision of the suicide vector, the deletion of an internal *toxT* fragment was confirmed via Sanger sequencing. In the Δ*toxT* derivatives, 165 internal amino acids (spanning the 41^st^ to 205^th^ amino acids) of ToxT are deleted.

### Construction of pBAD-toxT-His recombinant plasmids

Four 852-bp DNA fragments were PCR-amplified, each containing the following: the 828-bp *toxT* open reading frame (ORF) without the termination codon, with a GCC codon (Ala) inserted between the first and second amino acids to adjust the reading frame, an 18-bp sequence encoding six His residues at the C-terminus, and a TAA termination codon. The amplification was performed using the primer set toxT-Nco Ⅰ F: GGG CCA TGG CCA TTG GGA AAA AAT CTT TTC AAA and toxT-Sal Ⅰ HisR: CCG GTC GAC TTA *GTG GTG GTG GTG GTG GTG* TTT TTC TGC AAC TCC TGT CAA CA (His-tag sequence is shown in italics) on genomic DNA extracted from the O395 (*toxT-SY*) strain and its *toxT* allele derivatives: YJB001 (*toxT-SF*), EJK008 (*toxT-AY*), and EJK009 (*toxT-AF*). Each amplified DNA fragment was digested with NcoI and SalI and ligated into the corresponding restriction sites of the plasmid pBAD24 [42], resulting in the constructs pBAD-toxT-SY-His, pBAD-toxT-SF-His, pBAD-toxT-AY-His, and pBAD-toxT-AF-His.

The constructs were subsequently transformed into *E. coli* DH5α and *V. cholerae* DHL010 (N16961-Δ*toxT*). Expression of the His-tagged ToxT protein from the PBAD promoter was induced by adding L-arabinose to the bacterial culture at a final concentration of 0.2%. The expression of His-tagged ToxT was confirmed by Western blot analysis using an anti-His-tag (Cell Signaling, Danvers, MA).

### Construction of isogenic variants of *V. cholerae* strains that contain *toxT*-His

The DNA sequence of *tcpJ* is identical in both the classical and El Tor biotype strains of *V. cholerae* O1 serogroup. A 762-bp *tcpJ* fragment was PCR-amplified from the genomic DNA of the O395 strain using the primer pair TcpJ-Sal Ⅰ F: GGG GTC GAC ATG GAA TAC GTT TAC TTG and TcpJ-Sal Ⅰ R: GGG GTC GAC TCA CAT TAA ACG GAT TG. The amplified fragment was digested with SalI and inserted into the SalI site of the plasmids pBAD-toxT-SY-His, pBAD-toxT-SF-His, pBAD-toxT-AY-His, and pBAD-toxT-AF-His, generating pBAD-toxT-SY-His-tcpJ, pBAD-toxT-SF-His-tcpJ, pBAD-toxT-AY-His-tcpJ, and pBAD-toxT-AF-His-tcpJ, respectively. The orientation of *tcpJ* was verified by sequence analysis (Fig. S2).

Using these recombinant plasmids as templates, four 1,620-bp DNA fragments spanning from the initiation codon of His-tagged *toxT* to the termination codon of *tcpJ* were PCR-amplified with the primer pair pBAD-toxT-Xba Ⅰ F: GGC TCT AGA ATG GCC ATT GGG AAA and pBAD-tcpJ-Sac Ⅰ R: CGG GAG CTC TCA CAT TAA ACG GAT TG. These fragments were digested with XbaI and SacI and inserted into the corresponding sites of the suicide plasmid pCVD442, generating the constructs pCVD442-toxT-SY-His-tcpJ, pCVD442-toxT-SF-His-tcpJ, pCVD442-toxT-AY-His-tcpJ, and pCVD442-toxT-AF-His-tcpJ.

These recombinant plasmids were conjugally transferred into the corresponding isogenic *toxT* allele derivatives of the O395, N16961, and IB5230 strains to construct isogenic derivatives harboring His-tagged *toxT* through the allelic exchange method. Excision of the suicide vector and replacement of *toxT* with *toxT*-His were screened on sucrose-containing agar plates, and the DNA sequence of the His-tagged *toxT* was confirmed by Sanger sequencing (Fig. S2).

### CTX phage transduction

The transduction of CTXΦ into *V. cholerae* strains was carried out as previously described [23, 26]. Briefly, an isogenic derivative of *V. cholerae* N16961, referred to as PM20, in which the *ctxAB* genes were replaced with a kanamycin resistance cassette was used as the donor of CTXΦ. PM20 was cultured in LB medium supplemented with 20 ng/ml mitomycin C [18]. The resulting culture supernatant was then mixed with the O395 strain, which had been prepared under agglutinating conditions (30°C in LB medium, pH 6.5). After a 30-minute incubation to allow CTXΦ infection, the mixture was plated onto LB agar plates containing kanamycin (100 μg/ml). The presence of pCTX-1kan, the replicative form of the CTX phage genome, in the resulting transductants (O395:pCTX-1kan) was confirmed through Sanger sequencing of the intergenic region between the kanamycin resistance gene and *rstR*.

CTX-1kan virions, produced by culturing O395:pCTX-1kan in LB medium, were then used to transduce *V. cholerae* strains O395 (*toxT*-SY), O395-H (*toxT*-SY-His), EJK009 (*toxT*-AF), and EJK009-H (*toxT*-AF-His), all prepared under agglutinating conditions, as well as EJK009-H, which was prepared in LB medium at 37°C. Transduction efficiency was calculated by determining the ratio of transductants to the total number of recipient cells.

### SDS-PAGE and western blotting

Western blot analysis of His-tagged ToxT and TcpA was performed as described previously, with minor modifications [20]. Approximately 3 × 10^8^ (for His-tagged ToxT) or 1 × 10^7^ cells (for TcpA) were analyzed using 12% SDS-PAGE. The proteins separated by SDS-PAGE were transferred onto nitrocellulose membranes. Subsequently, western blot analysis was performed using anti-His (Cell Signaling, Danvers, MA) and anti-TcpA (a generous gift from Dr. W. F. Wade, Dartmouth University, Hanover, NH, USA [43]), with HRP-linked goat anti-rabbit IgG, (7074S, Cell Signaling, Danvers, MA, USA) as the secondary antibody.

### RT-PCR

A real-time qRT-PCR assay was performed as previously described, with some modifications [13]. Total RNA was extracted from approximately 2 × 10^9^ cells at each time point using the Nextractor-48N® nucleic acid extractor system (Genolution, Seoul, Korea), followed by treatment with RNase-free DNase 1. cDNA was synthesized with 1 μg of RNA using the Thermo Scientific RevertAid First Strand cDNA synthesis kit (ThermoFisher). The concentrations of RNA and cDNA were determined using NanoDrop 2000 (ThermoFisher). Real-time PCR was carried out using 100 ng of synthesized cDNA and specific primers (Table S3) with a CFX96RT-PCR system and iQ^TM^ SYBR® Green Supermix (BioRad). The PCR conditions were 95°C for 2 min, followed by 35 cycles of 95°C for 10 s, 60°C for 30 s, and 72°C for 30 s. Each run concluded with a melting-curve ranging from 55°C to 95°C to validate specific amplification. Expression values were normalized to the housekeeping gene *gyrA* as described previously [13, 34, 44]. The relative expression levels of *toxT*, *tcpA*, and *ctxAB* were calculated using the 2^-ΔΔCT^ method [34]. Data were collected from three independent experiments.

## Author contributions

E.J.K. and D.W.K. conceived the project. E.J.K, J.J., D.L., S. C., and H. K. prepared samples performed experiments, E.J.K and D.W.K wrote the first draft of the paper and all authors reviewed, edited, and commented on the draft.

## Additional information

Correspondence and requests for materials should be addressed to Dong Wook Kim or Eun Jin Kim

## Supporting Information

**Table S1.** CTXΦ Transduction efficiency of selected *V. cholerae* strains.

|  | **O395**<br>(*toxT*-SY) | **O395-H**<br>(*toxT*-SY-His) | **EJK009**<br>(*toxT*-AF) | **EJK009-H**<br>(*toxT*-AF-His) | **EJK009-H**<br>(37°C) |
| --- | --- | --- | --- | --- | --- |
| No. of<br>transductants /<br>recipient cells | $\frac{5.5 \times 10^7}{3.0 \times 10^8}$ | $\frac{4.3 \times 10^7}{2.7 \times 10^8}$ | $\frac{7.3 \times 10^7}{3.6 \times 10^8}$ | $\frac{3.6 \times 10^7}{1.8 \times 10^8}$ | $\frac{1.4 \times 10^4}{2.8 \times 10^8}$ |
| Transduction<br>Efficiency | 18.3% | 15.9% | 20.3% | 20.0% | 0.005% |

**Table S2.** *V. cholerae* strains and their isogenic derivatives.

| Strains | *toxT* genotype | Genome information and references | Strains | *toxT* genotype | Genome information and references |
| --- | --- | --- | --- | --- | --- |
| Classical biotype |  |  | El Tor Biotype |  |  |
| O395 | O395-*toxT*-SY | CP000626/CP000627 [3] | N16961 | N16961-*toxT*-SY | AE003852/AE003853 [45] |
| O395-H | O395-*toxT*-SY-His | This study | N16961-H | N16961-*toxT*-SY-His | This study |
| YJB001 | O395-*toxT*-SF | [21] | YJB003 | N16961-*toxT*-SF | [21] |
| YJB001-H | O395-*toxT*-SF-His | This study | YJB003-H | N16961-*toxT*-SF-His | This study |
| EJK008 | O395-*toxT*-AY | [20] | DHL008 | N16961-*toxT*-AY | [20] |
| EJK008-H | O395-*toxT*-AY-His | This study | DHL008-H | N16961-*toxT*-AY-His | This study |
| EJK009 | O395-*toxT*-AF | [20] | DHL009 | N16961-*toxT*-AF | [20] |
| EJK009-H | O395-*toxT*-AF-His | This study | DHL009-H | N16961-*toxT*-AF-His | This study |
| EJK010 | O395-Δ*toxT* | [20] | DHL010 | N16961-Δ*toxT* | [20] |
| 569B | 569B-*toxT*-AY | DADXPZ010000000 |  |  |  |
| EJK007 | 569B-*toxT*-AF | [22] | IB5230 | IB5230-*toxT*-SY | AELH00000000.1 [27] |
|  |  |  | IB5230-H | IB5230-*toxT*-SY-His | This study |
|  |  |  | YJB020 | IB5230-*toxT*-SF | [21] |
|  |  |  | YJB020-H | IB5230-*toxT*-SF-His | This study |
|  |  |  | DHL020 | IB5230-*toxT*-AY | [20] |
|  |  |  | DHL020-H | IB5230-*toxT*-AY-His | This study |
|  |  |  | DHL021 | IB5230-*toxT*-AF | [20] |
|  |  |  | DHL021-H | IB5230-*toxT*-AF-His | This study |
| | | | DHL022 | IB5230-Δ*toxT* | [20] |

*toxT*-SY: *toxT* allele with Ser at position 65 and Tyr at position 139.
*toxT*-SF: *toxT* allele with Ser at position 65 and Phe at position 139.
*toxT*-AY: *toxT* allele with Ala at position 65 and Tyr at position 139.
*toxT*-AF: *toxT* allele with Ala at position 65 and Phe at position 139.

**Table S3.**
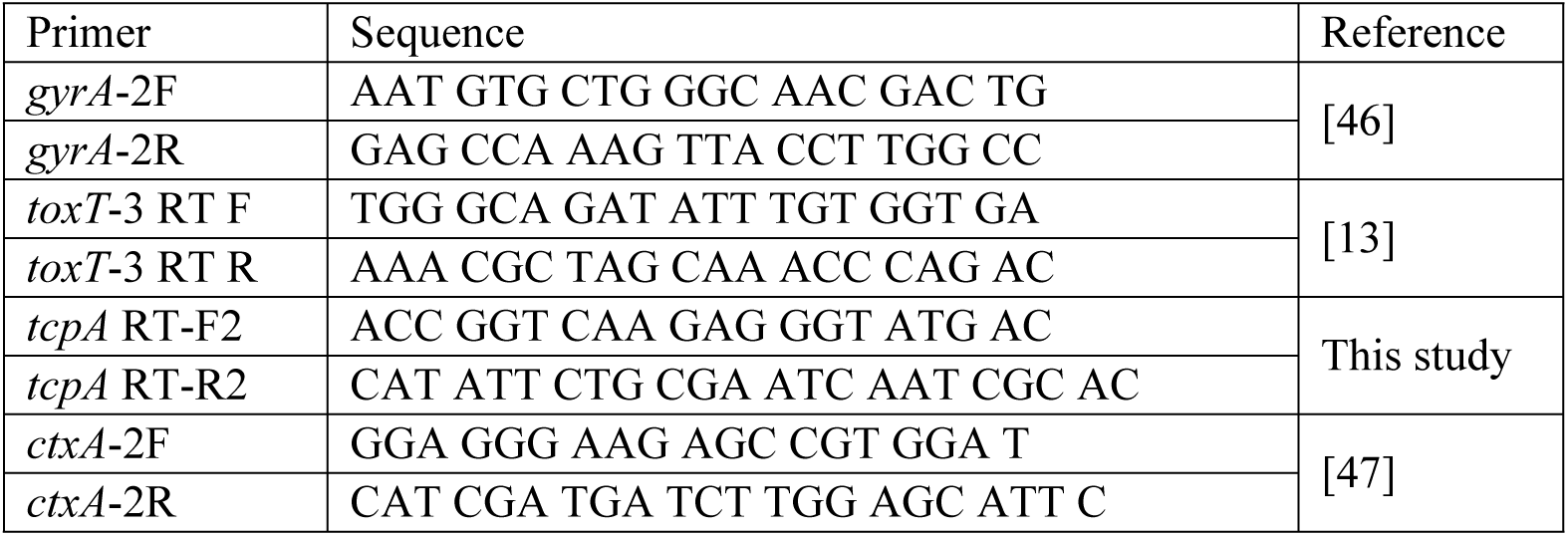
Oligonucleotide sequences for qRT-PCR.

**Figure S1.**
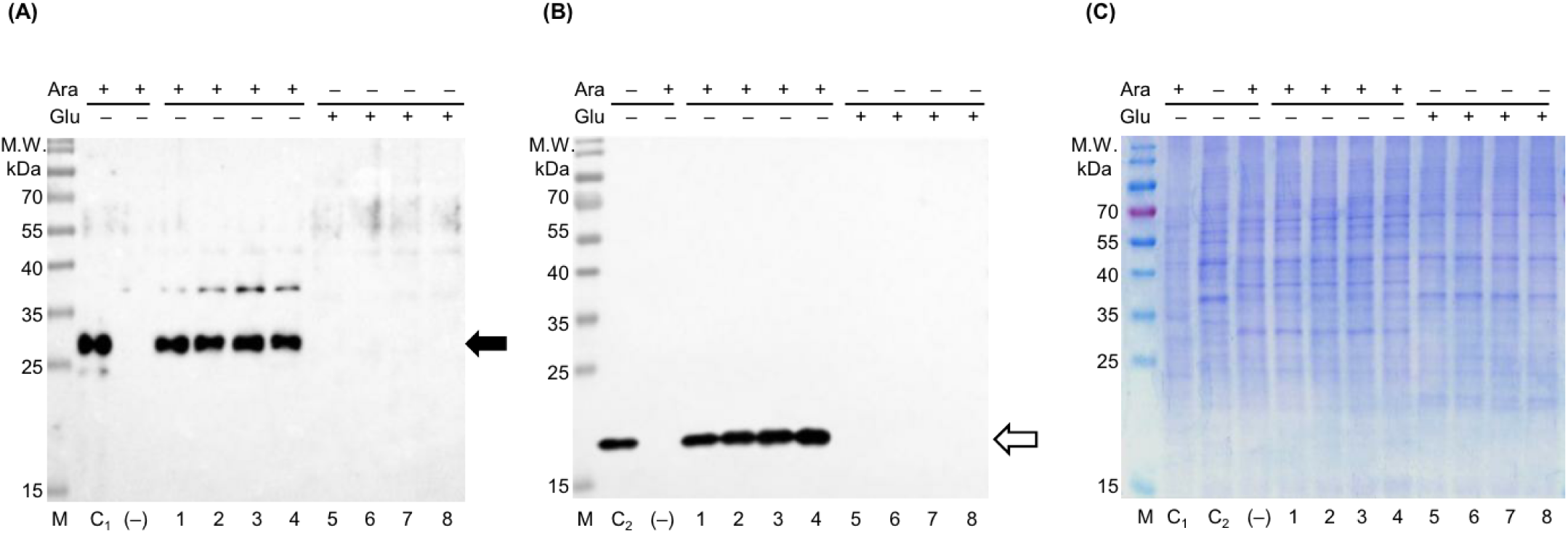
Expression of ToxT and TcpA in a *toxT*-deleted isogenic derivative of N16961 that harbors the pBAD plasmid containing the *toxT* allele. ToxT, driven by the P_BAD_ promoter, and TcpA were analyzed by Western blotting using anti-His-tag and anti-TcpA antibodies. The bacteria were cultured in LB medium at 30°C with 0.2% arabinose or 0.5% glucose. (A) Western blot analysis using anti-His-tag antibodies. His-tagged ToxT is indicated by a black arrow. Lane C_1_: *E. coli* DH5α harboring pBAD-*toxT*-SY cultured in LB medium with 0.2% arabinose. Lane (–): DHL010 (N16961-Δ*toxT*) harboring the blank pBAD24 plasmid, cultured in LB medium with 0.2% arabinose. DHL010 containing pBAD-*toxT-*SY (Lanes 1 and 5), pBAD-*toxT*-SF (Lanes 2 and 6), pBAD-*toxT-*AY (Lanes 3 and 7), and pBAD-*toxT*-AF (Lanes 4 and 8) were cultured in LB medium with 0.2% arabinose (lanes 1**–**4) or 0.5% glucose (lanes 5**–**8). (B) Western blotting analysis using anti-TcpA antibodies. Lane C_2_: O395 cultured in LB medium (pH 6.5) at 30°C. Other lanes are treated as described in Panel (A). The white arrow indicates TcpA. (C): Coomassie Brilliant Blue staining of SDS-PAGE for all bacterial samples. Approximately 10^7^ cells were analyzed in each lane.

**Figure S2.**
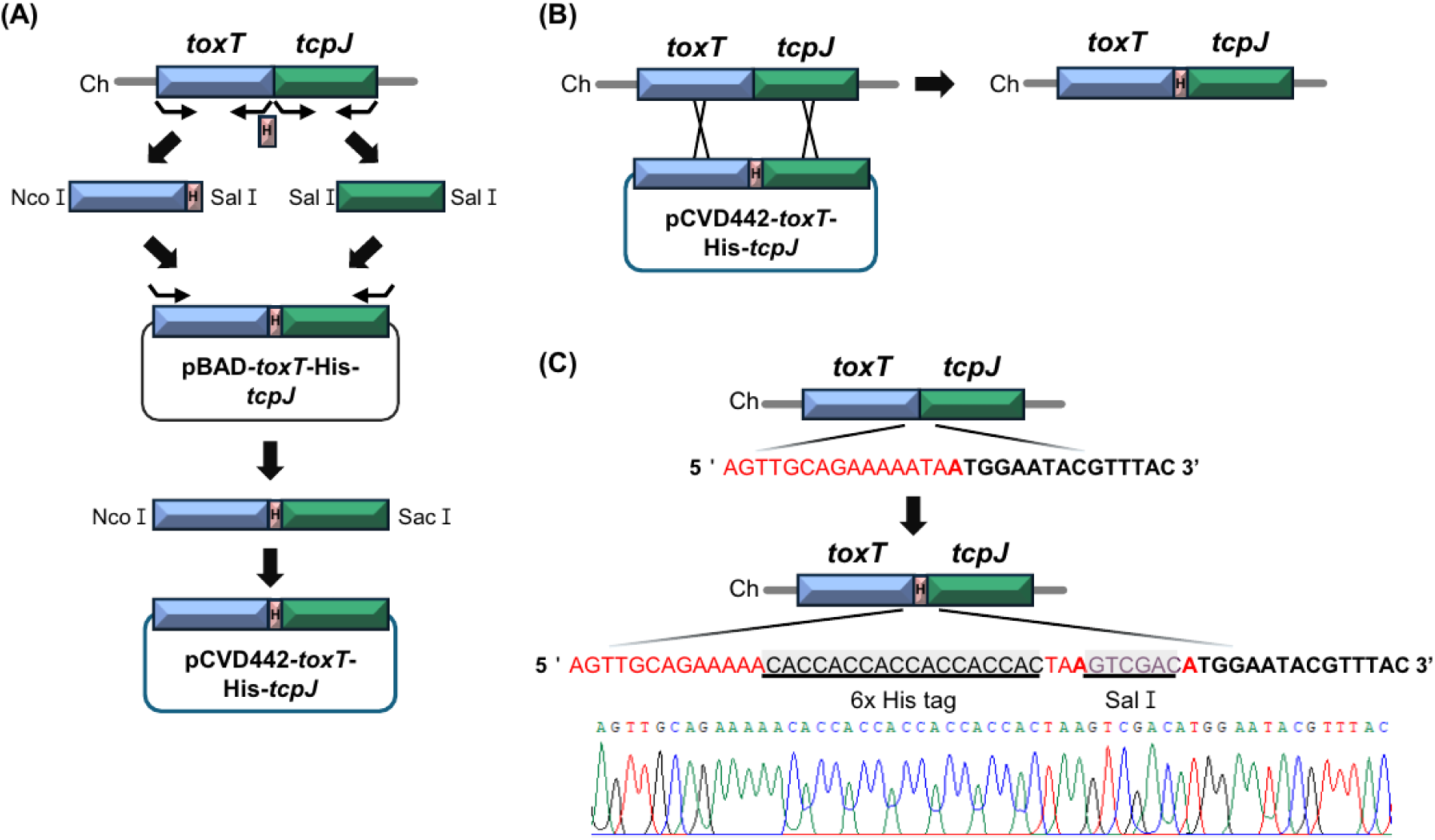
Construction of isogenic derivatives of *V. cholerae* strains that harbor His-tagged *toxT*. (A) Construction of a set of suicide plasmids, pCVD442-toxT-His-tcpJs, containing different *toxT* alleles (described in detail in the Materials and Methods section). (B) Allelic exchange of native *toxT* with His-tagged *toxT*. Each of the native *toxT* alleles was replaced by its corresponding His-tagged *toxT* allele. (C) DNA sequence comparison between the junctions of native *toxT*-*tcpJ* and His-tagged *toxT*-*tcpJ*.

**Figure S3.**
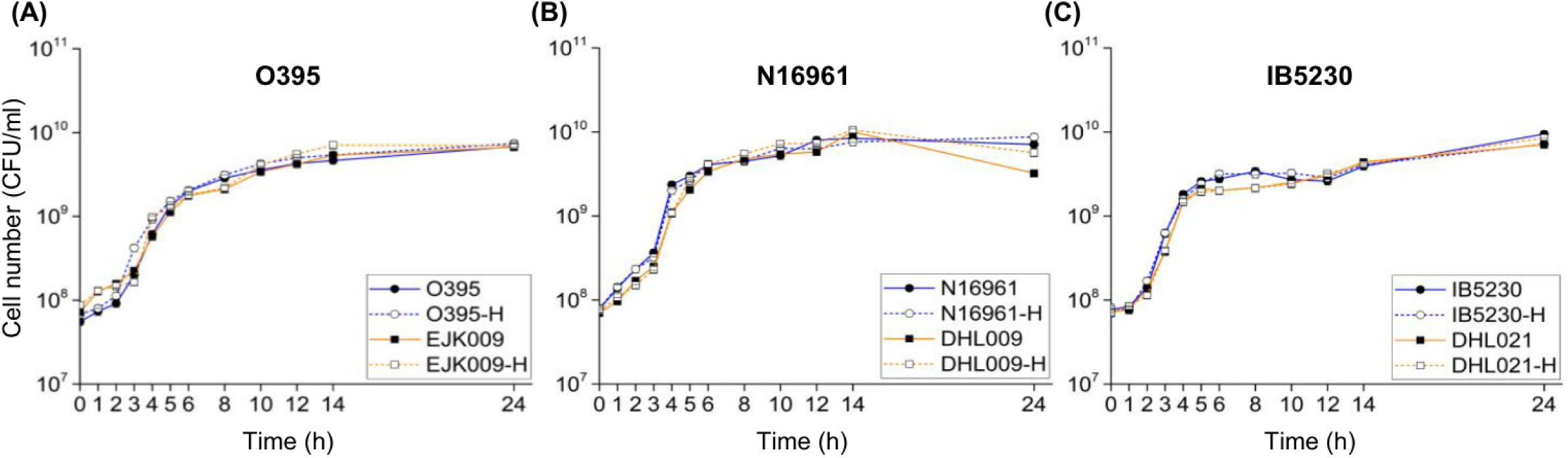
The growth curves of O395, N16961, and IB5230 and their *toxT* allele isogenic derivatives. (A) Growth curves of O395 and its isogenic derivatives [O395 (*toxT*-SY): blue solid line, O395-H (*toxT*-SY-His): blue dotted line, EJK009 (*toxT*-AF): orange solid line, and EJK009-H (*toxT*-AF-His): orange dotted line] represented by colony-forming units per milliliter (CFU/ml) over time (hours). (B) Growth curves of N16961 and isogenic derivatives [N16961 (*toxT*-SY): blue solid line, N16961-H (*toxT*-SY-His): blue dotted line, DHL009 (*toxT*-AF): orange solid line, and DHL009-H (*toxT*-AF-His): orange dotted line]. (C) Growth curves of IB5230 and its isogenic derivatives [IB5230 (*toxT*-SY): blue solid line, IB5230-H (*toxT*-SY-His): blue dotted line, DHL021 (*toxT*-AF): orange solid line, and DHL021-H (*toxT*-AF-His): orange dotted line]. Bacteria were cultured in LB medium at 30°C. Standard deviation of cell numbers was omitted for clarity.

**Figure S4.**
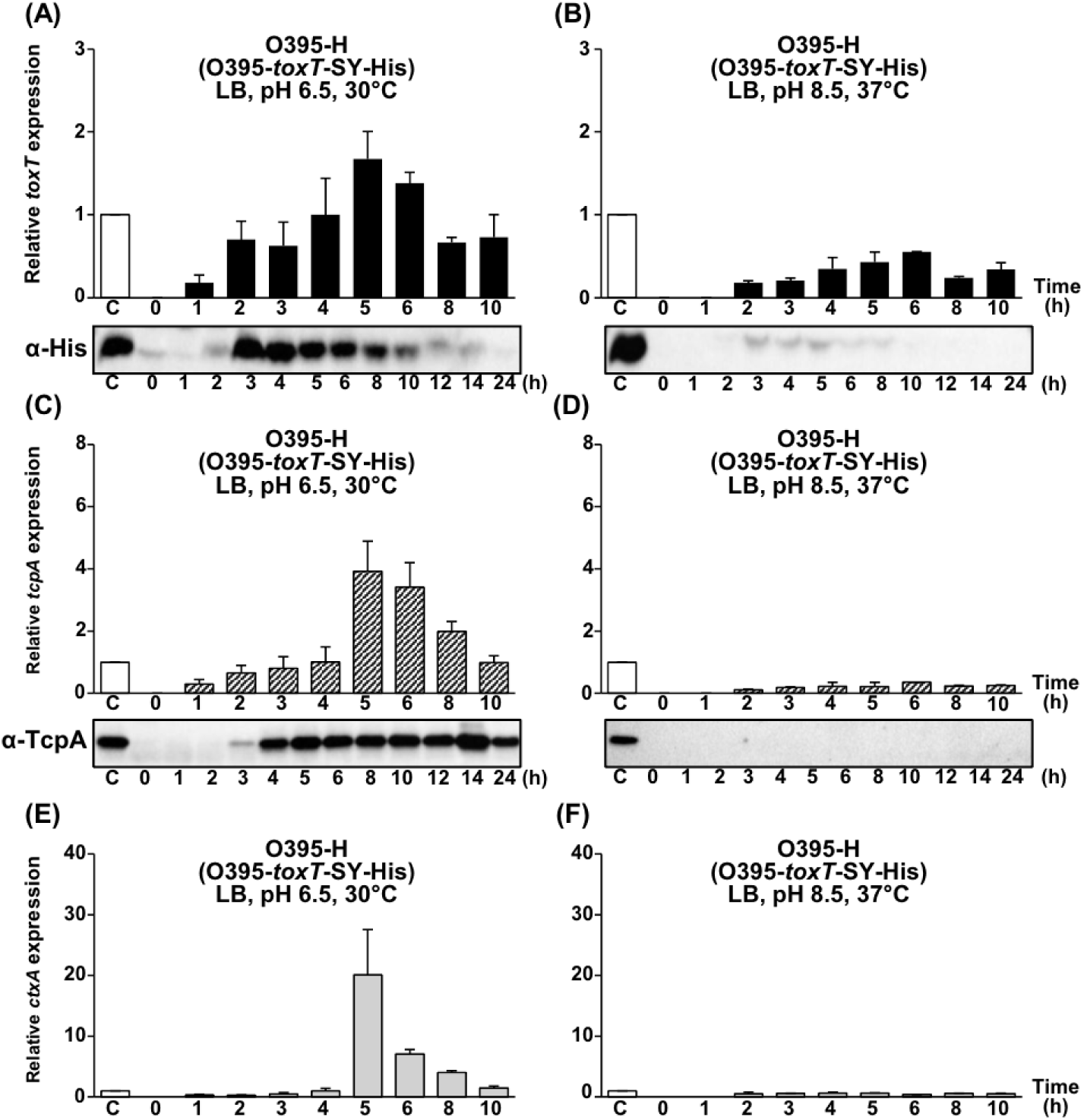
Expression of His-tagged *toxT*, *tcpA*, and *ctxAB* in O395-H (O395-*toxT*-SY-His). (A, B) qRT-PCR analysis of His-tagged *toxT* mRNA and Western blot detection of ToxT using anti-His-tag antibodies in O395-H cultured in (A) LB medium (pH 6.5) at 30°C and (B) LB medium (pH 8.5) at 37°C. (C, D) qRT-PCR analysis of *tcpA* mRNA and Western blot detection of TcpA using anti-TcpA antibodies in O395-H cultured in (C) LB (pH 6.5) at 30°C and (D) LB (pH 8.5) at 37°C. (E, F) qRT-PCR analysis of *ctxAB* mRNA in O395-H cultured in (E) LB (pH 6.5) at 30°C and (F) LB (pH 8.5) at 37°C. Expression values were normalized to the housekeeping gene *gyrA*. The relative expression levels of His-tagged *toxT* (black bar), *tcpA* (diagonal bar), and *ctxAB* (gray bar) at each time point are presented, with their expression levels in the 4-hour culture of O395-H grown in LB (pH 6.5) at 30°C set to 1 (lane C, white bar).

**Figure S5.**
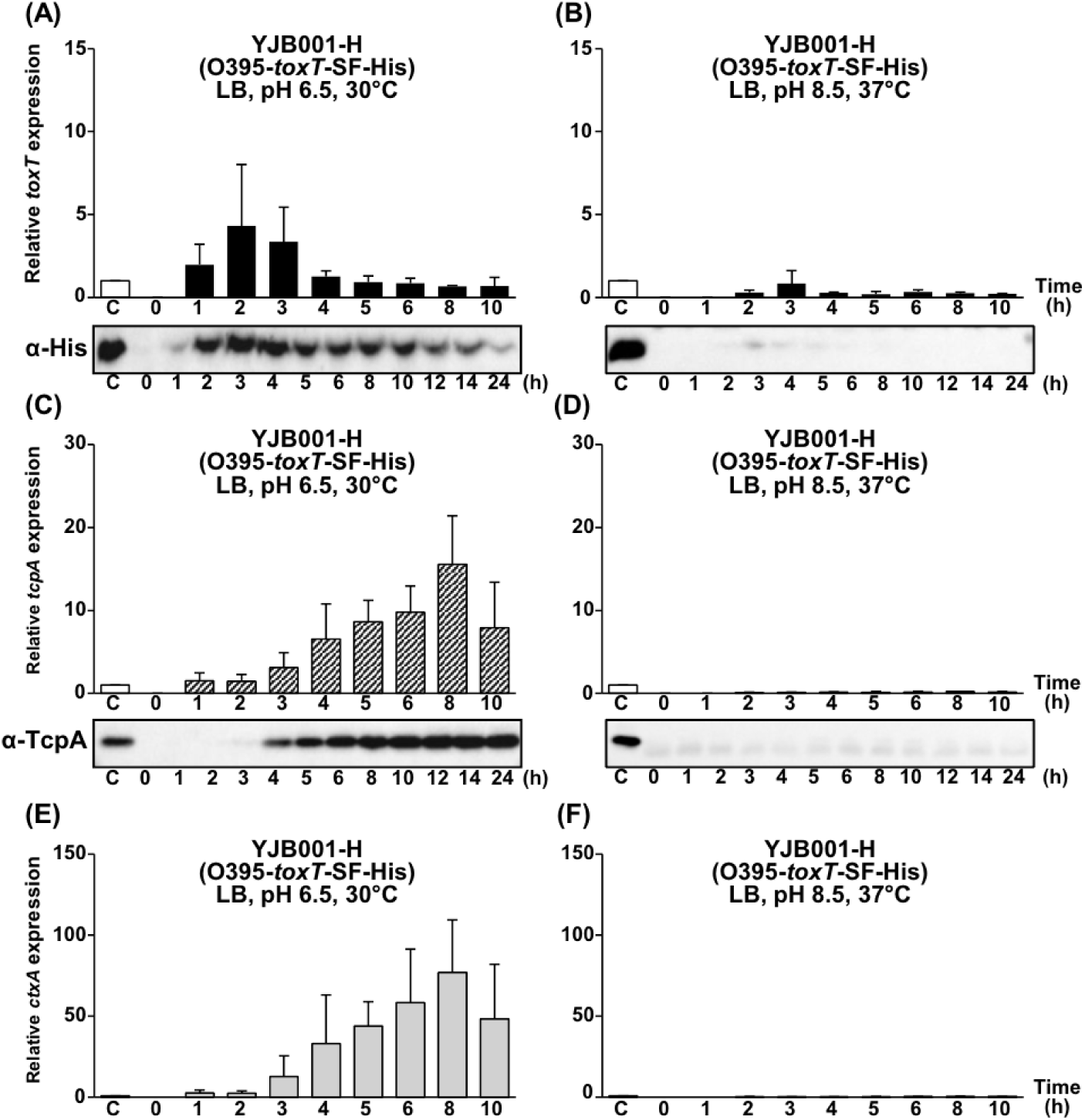
Expression of His-tagged *toxT*, *tcpA*, and *ctxAB* in YJB001-H (O395-*toxT*-SF-His). (A, B) qRT-PCR analysis of His-tagged *toxT* mRNA and Western blot detection of ToxT using anti-His-tag antibodies in YJB001-H cultured in (A) LB medium (pH 6.5) at 30°C and (B) LB (pH 8.5) at 37°C. (C, D) qRT-PCR analysis of *tcpA* mRNA and Western blot detection of TcpA using anti-TcpA antibodies in YJB001-H cultured in (C) LB (pH 6.5) at 30°C and (D) LB (pH 8.5) at 37°C. (E, F) qRT-PCR analysis of *ctxAB* mRNA in YJB001-H cultured in (E) LB (pH 6.5) at 30°C and (F) LB (pH 8.5) at 37°C. Expression values were normalized to the housekeeping gene *gyrA*. The relative expression levels of His-tagged *toxT* (black bar), *tcpA* (diagonal bar), and *ctxAB* (gray bar) at each time point are presented, with their expression levels in the 4-hour culture of O395-H grown in LB (pH 6.5) at 30°C set to 1 (lane C, white bar).

**Figure S6.**
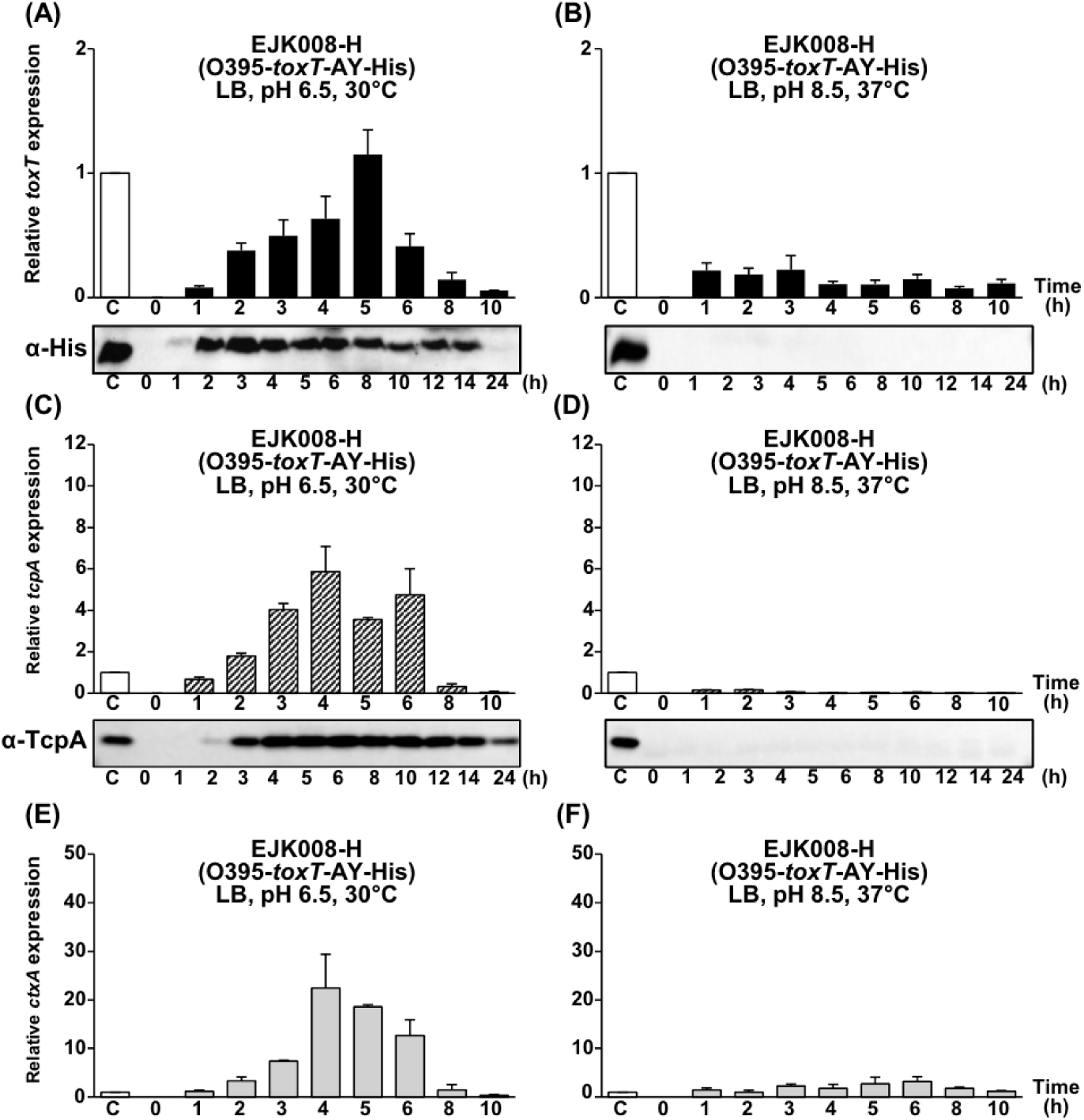
Expression of His-tagged *toxT*, *tcpA*, and *ctxAB* in EJK008-H (O395-*toxT*-AY-His). (A, B) qRT-PCR analysis of His-tagged *toxT* mRNA and Western blot detection of ToxT using anti-His-tag antibodies in EJK008-H cultured in (A) LB medium (pH 6.5) at 30°C and (B) LB (pH 8.5) at 37°C. (C, D) qRT-PCR analysis of *tcpA* mRNA and Western blot detection of TcpA using anti-TcpA antibodies in EJK008-H cultured in (C) LB (pH 6.5) at 30°C and (D) LB (pH 8.5) at 37°C. (E, F) qRT-PCR analysis of *ctxAB* mRNA in EJK008-H cultured in (E) LB (pH 6.5) at 30°C and (F) LB (pH 8.5) at 37°C. Expression values were normalized to the housekeeping gene *gyrA*. The relative expression levels of His-tagged *toxT* (black bar), *tcpA* (diagonal bar), and *ctxAB* (gray bar) at each time point are presented, with their expression levels in the 4-hour culture of O395-H grown in LB (pH 6.5) at 30°C set to 1 (lane C, white bar).

**Figure S7.**
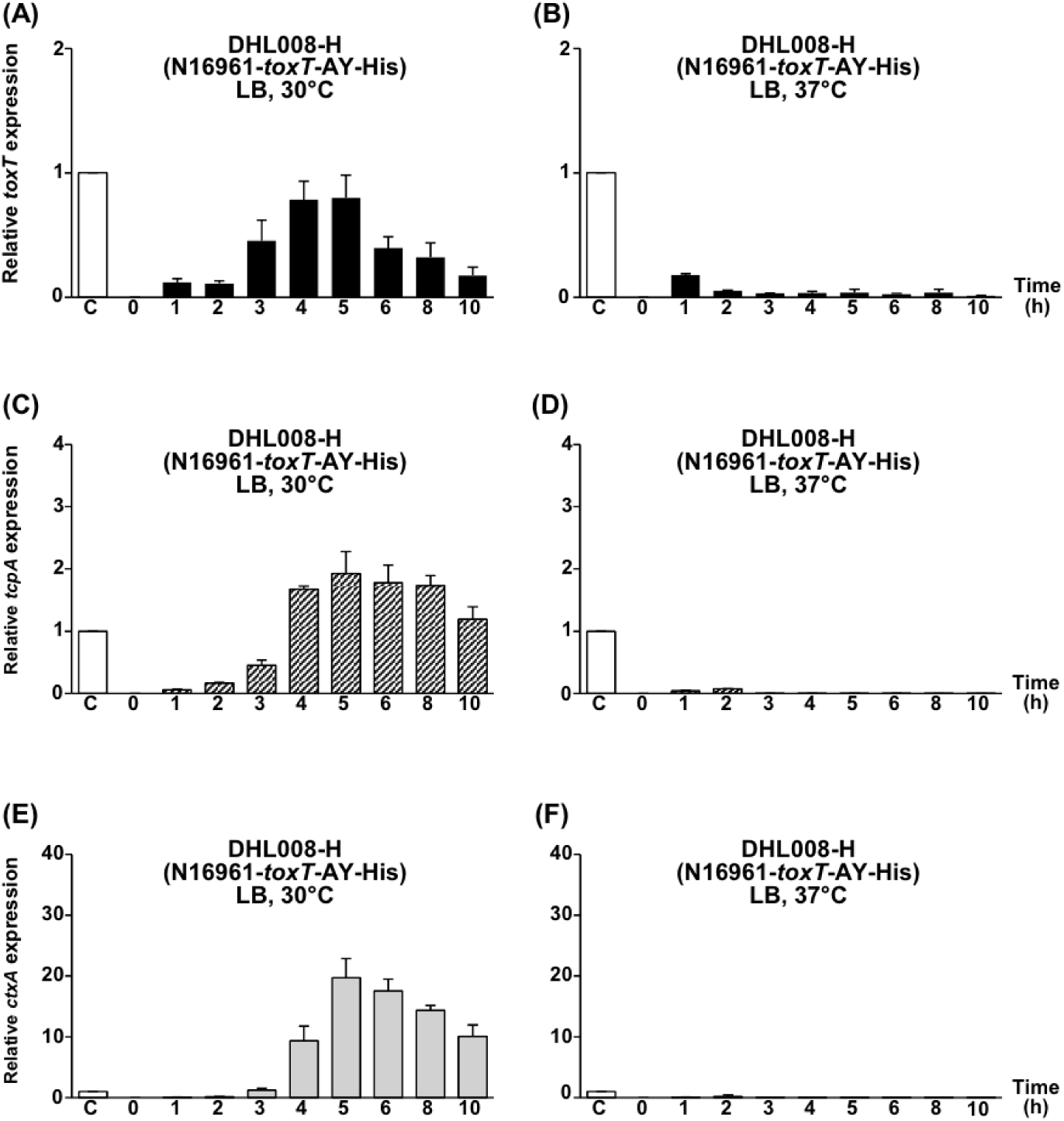
Expression of His-tagged *toxT*, *tcpA*, and *ctxAB* in DHL008-H (N16961-*toxT*-AY-His). (A, B) qRT-PCR analysis of *toxT*-SF-His mRNA in DHL008-H cultured in LB at (A) 30°C and (B) 37°C. (C, D) qRT-PCR analysis of *tcpA* mRNA in DHL008-H cultured in LB at (C) 30°C and (D) 37°C. (E, F) qRT-PCR analysis of *ctxAB* mRNA in DHL008-H cultured in LB at (E) 30°C and (F) 37°C. Expression values were normalized to the housekeeping gene *gyrA*. The relative expression levels of His-tagged *toxT* (black bar), *tcpA* (diagonal bar), and *ctxAB* (gray bar) at each time point are presented, with their expression levels in the 4-hour culture of O395-H grown in LB (pH 6.5) at 30°C set to 1 (lane C, white bar).

**Figure S8.**
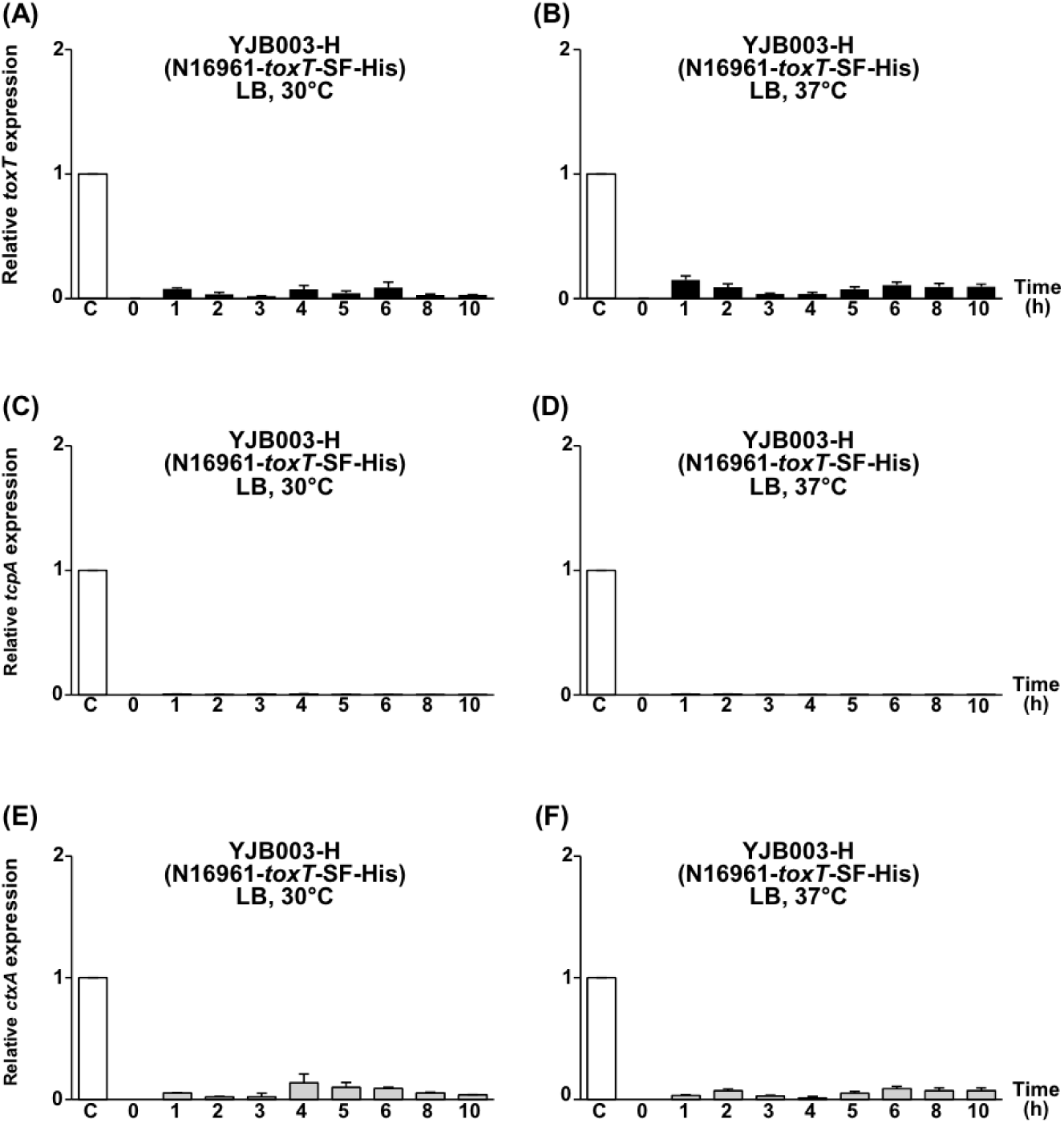
Expression of His-tagged *toxT*, *tcpA*, and *ctxAB* in YJB003-H (N16961-*toxT*-SF-His). (A, B) qRT-PCR analysis of *toxT*-SF-His mRNA in YJB003-H cultured in LB at (A) 30°C and (B) 37°C. (C, D) qRT-PCR analysis of *tcpA* mRNA in YJB003-H cultured in LB at (C) 30°C and (D) 37°C. (E, F) qRT-PCR analysis of *ctxAB* mRNA in YJB003-H cultured in LB at (E) 30°C and (F) 37°C. Expression values were normalized to the housekeeping gene *gyrA*. The relative expression levels of His-tagged *toxT* (black bar), *tcpA* (diagonal bar), and *ctxAB* (gray bar) at each time point are presented, with their expression levels in the 4-hour culture of O395-H grown in LB (pH 6.5) at 30°C set to 1 (lane C, white bar).

**Figure S9.**
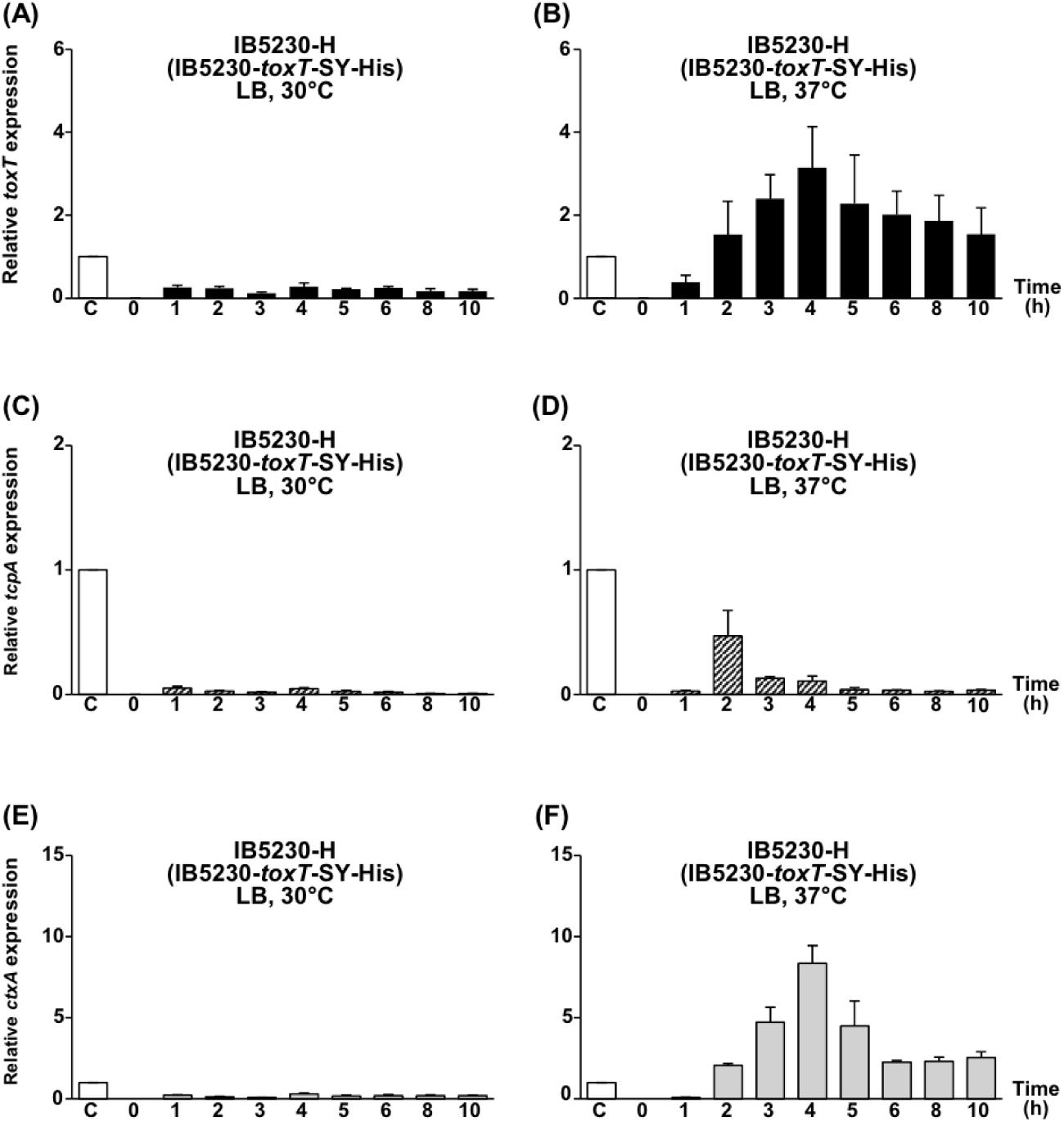
Expression of His-tagged *toxT*, *tcpA*, and *ctxAB* in IB5230-H (IB5230-*toxT*-SY-His). (A, B) qRT-PCR analysis of *toxT*-SF-His mRNA in IB5230-H cultured in LB at (A) 30°C and (B) 37°C. (C, D) qRT-PCR analysis of *tcpA* mRNA in IB5230-H cultured in LB at (C) 30°C and (D) 37°C. (E, F) qRT-PCR analysis of *ctxAB* mRNA in IB5230-H cultured in LB at (E) 30°C and (F) 37°C. Expression values were normalized to the housekeeping gene *gyrA*. The relative expression levels of His-tagged *toxT* (black bar), *tcpA* (diagonal bar), and *ctxAB* (gray bar) at each time point are presented, with their expression levels in the 4-hour culture of O395-H grown in LB (pH 6.5) at 30°C set to 1 (lane C, white bar).

**Figure S10.**
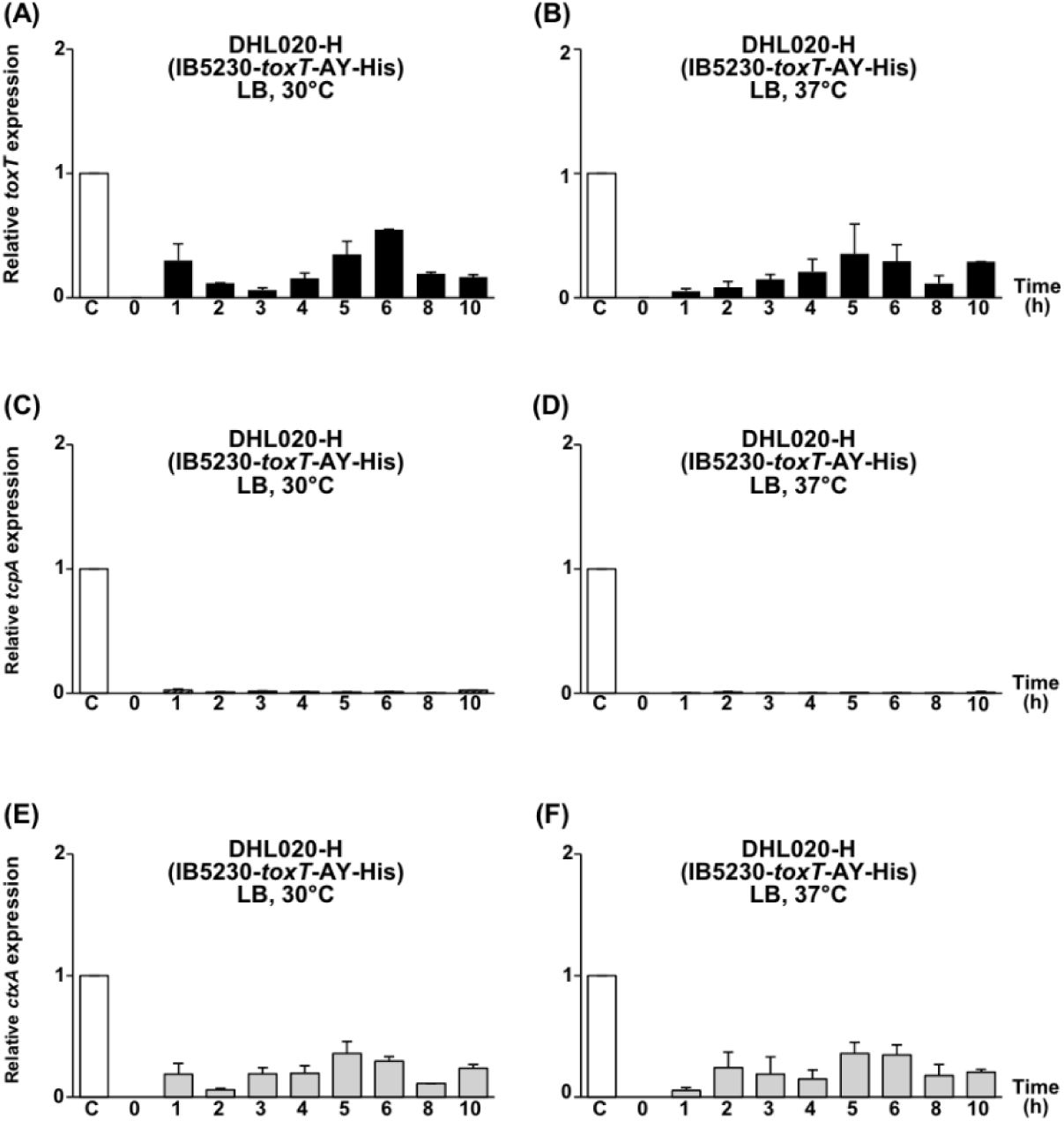
Expression of His-tagged *toxT*, *tcpA*, and *ctxAB* in DHL020-H (IB5230-*toxT*-AY-His). (A, B) qRT-PCR analysis of *toxT*-SF-His mRNA in DHL020-H cultured in LB at (A) 30°C and (B) 37°C. (C, D) qRT-PCR analysis of *tcpA* mRNA in DHL020-H cultured in LB at (C) 30°C and (D) 37°C. (E, F) qRT-PCR analysis of *ctxAB* mRNA in DHL020-H cultured in LB at (E) 30°C and (F) 37°C. Expression values were normalized to the housekeeping gene *gyrA*. The relative expression levels of His-tagged *toxT* (black bar), *tcpA* (diagonal bar), and *ctxAB* (gray bar) at each time point are presented, with their expression levels in the 4-hour culture of O395-H grown in LB (pH 6.5) at 30°C set to 1 (lane C, white bar).

**Fig. S11.**
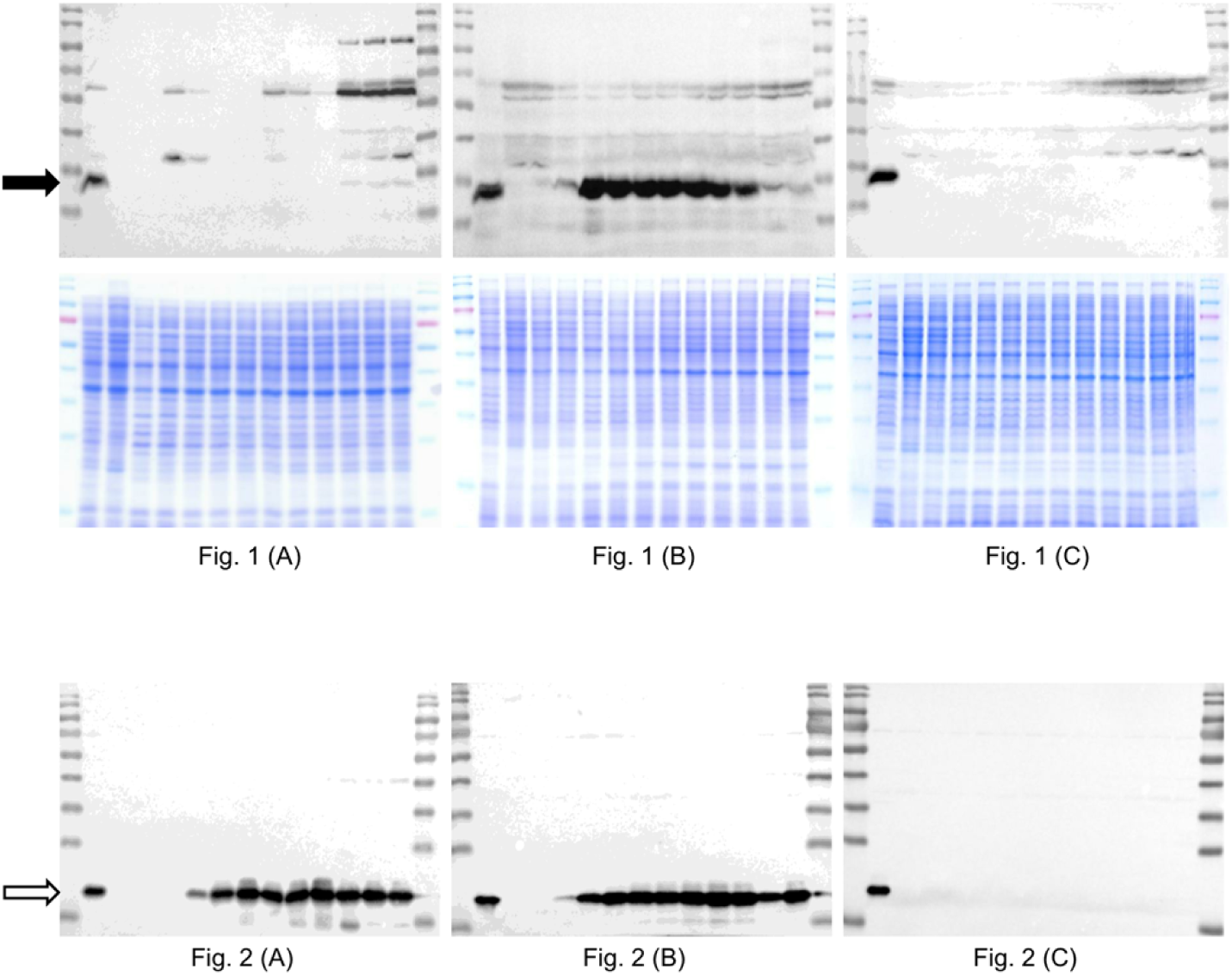
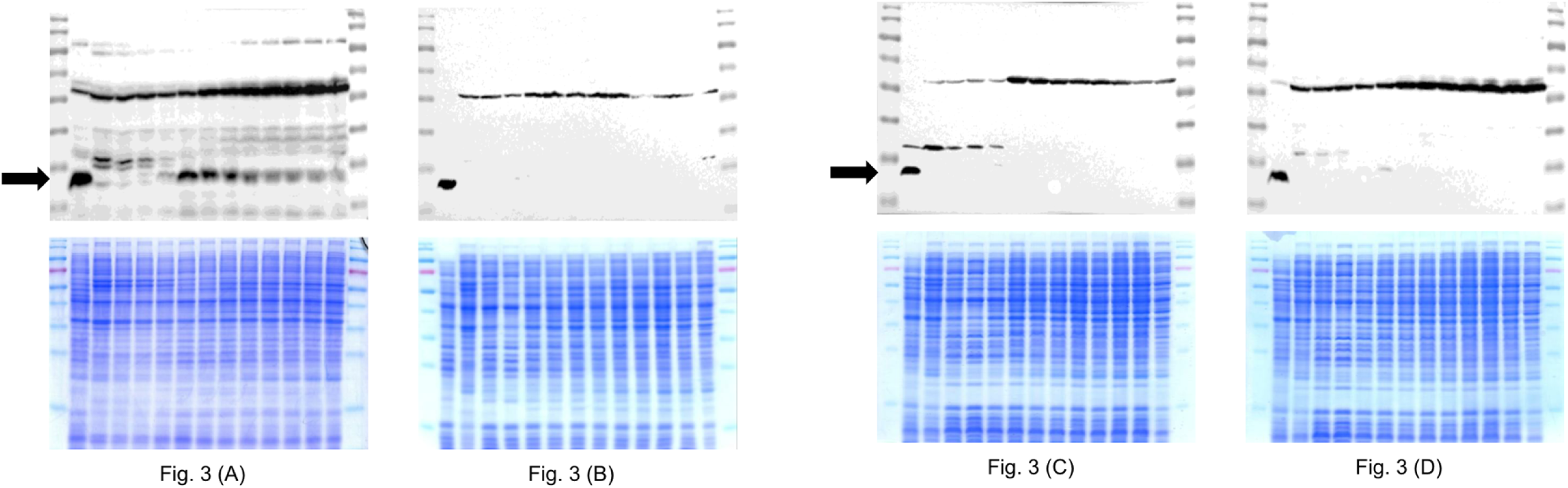
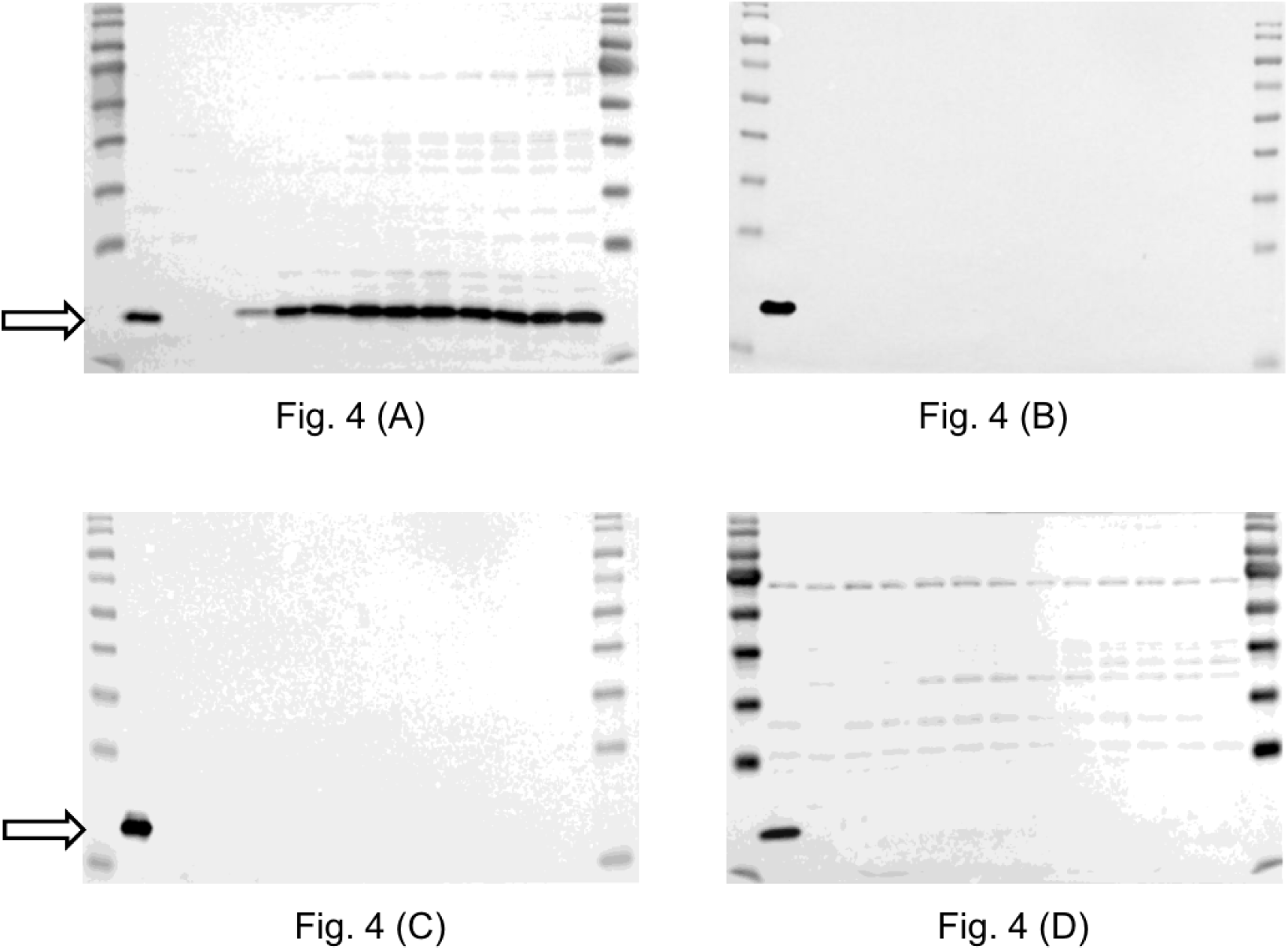
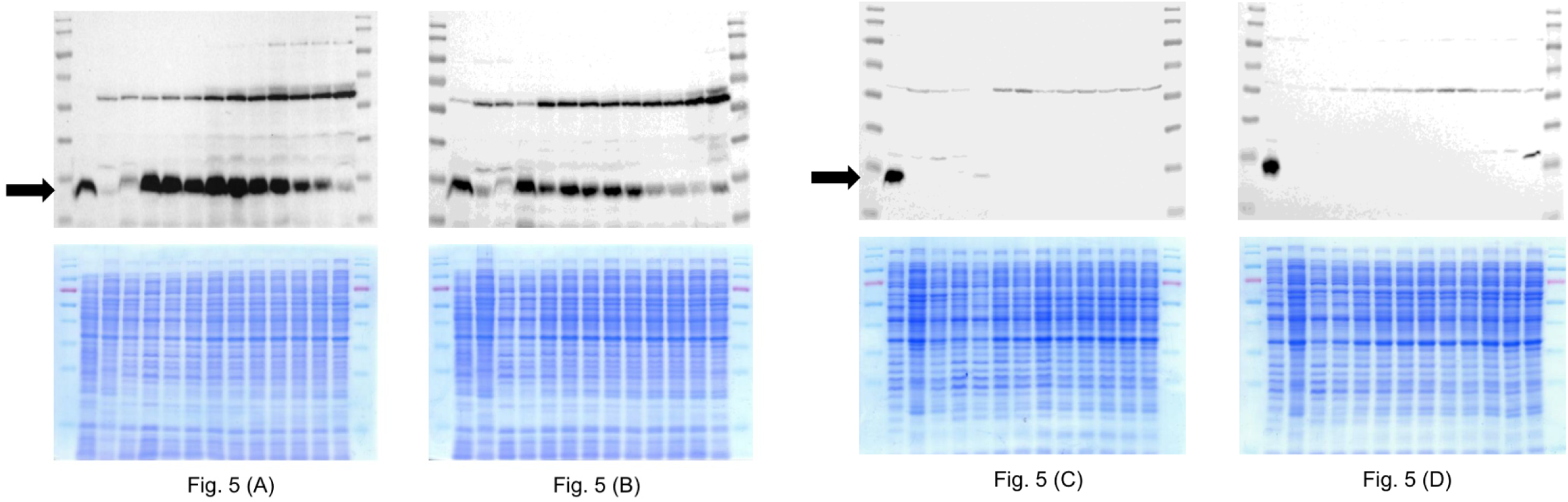
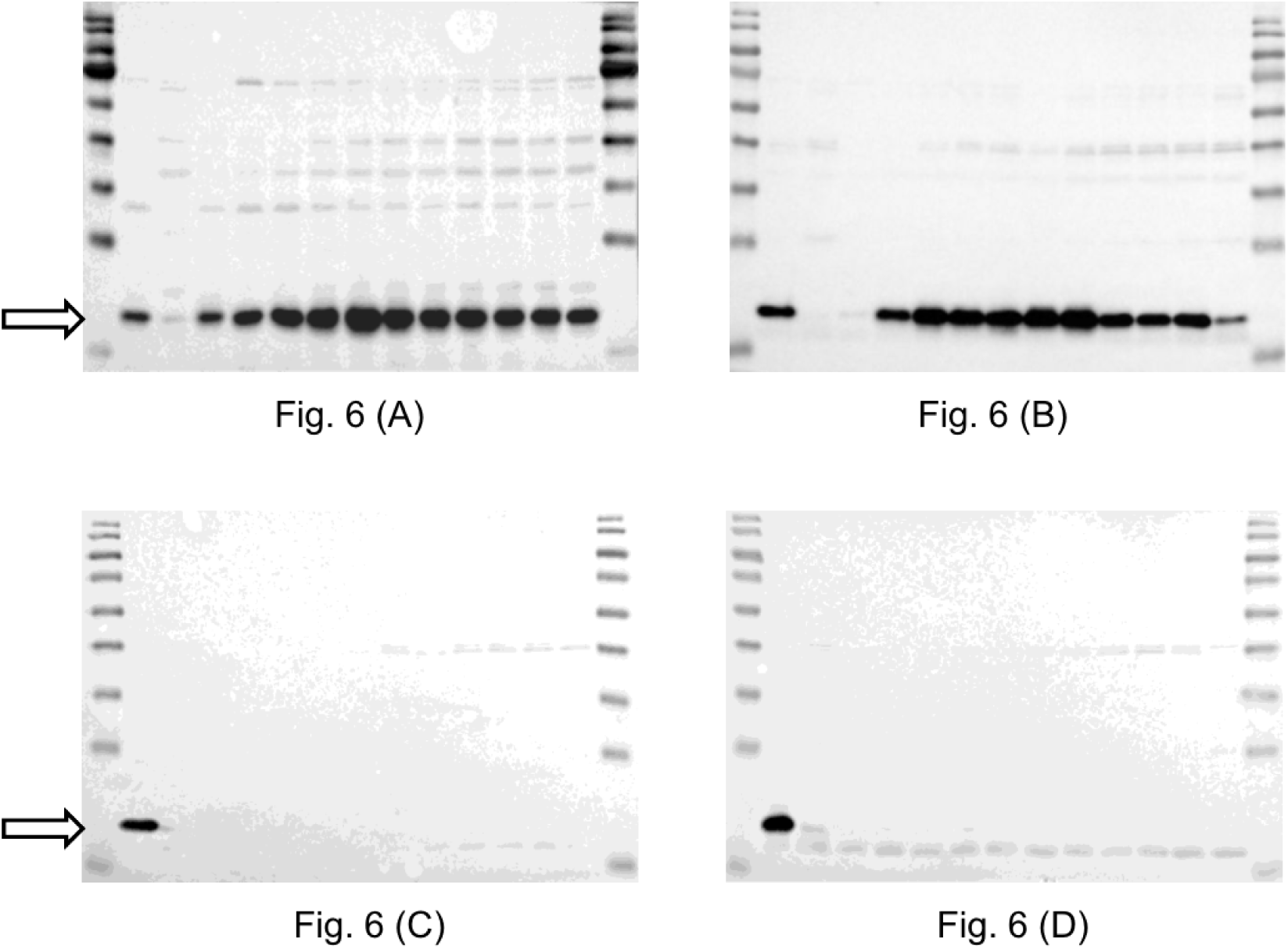
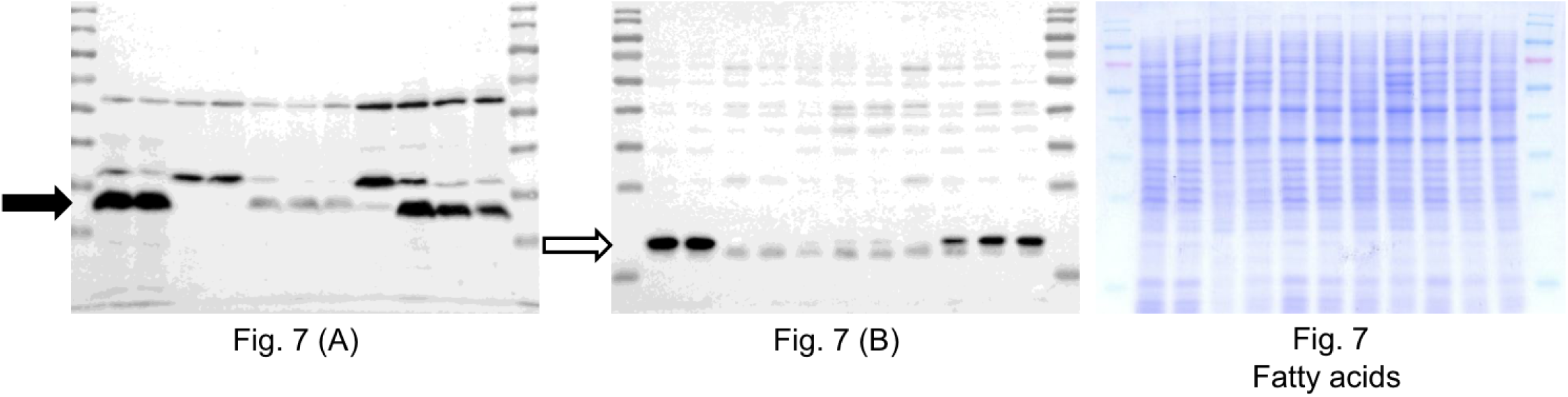
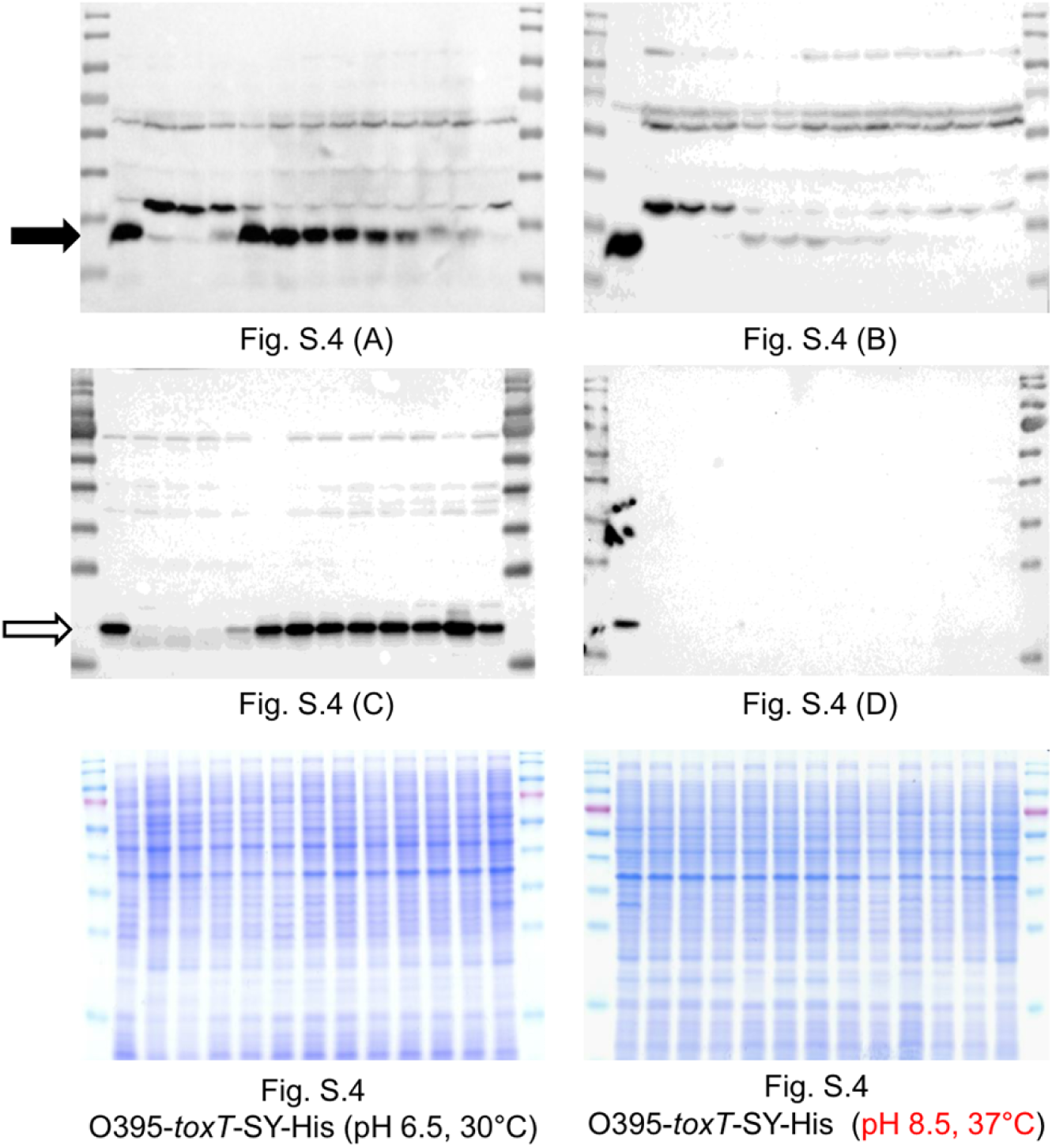
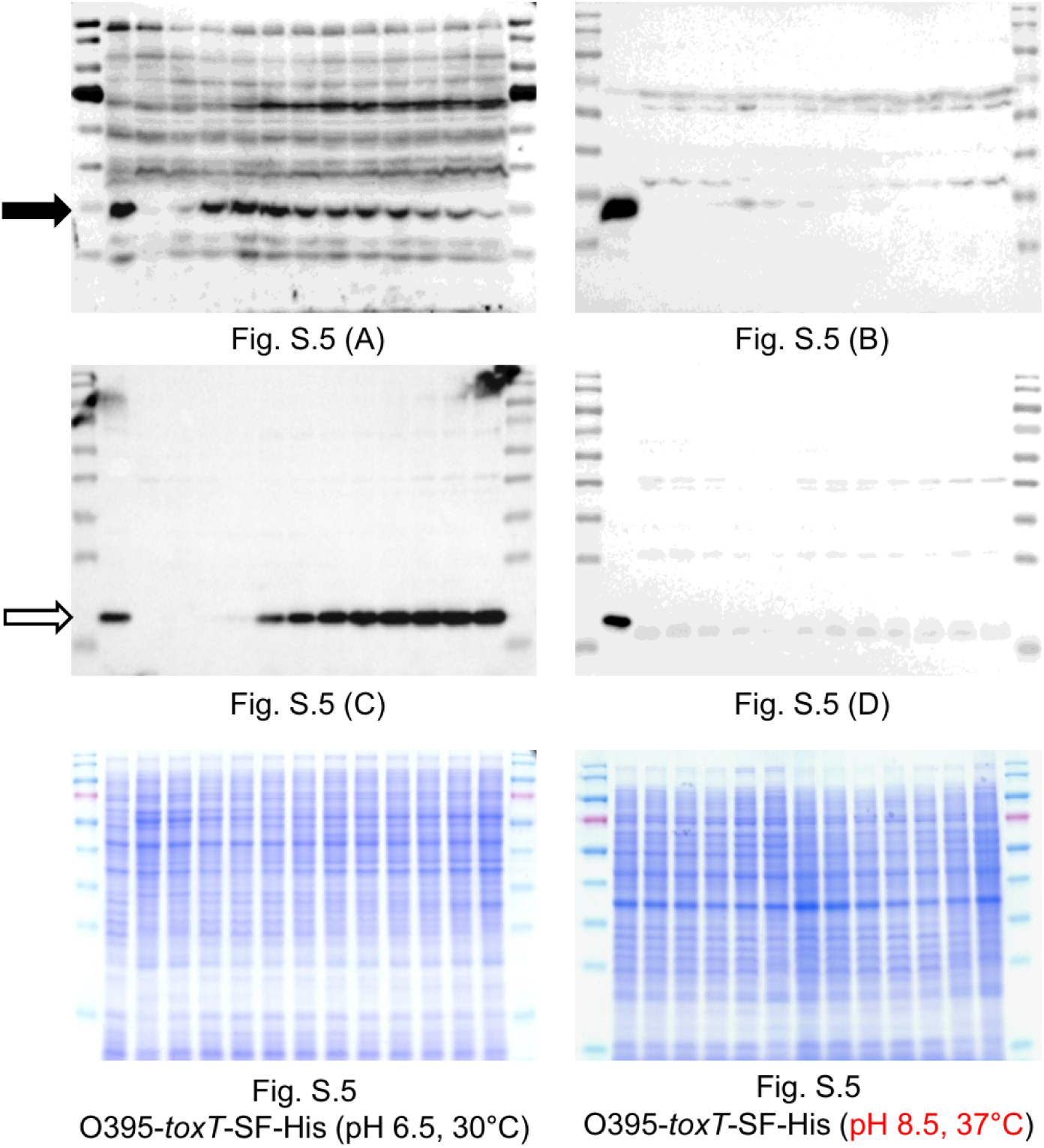
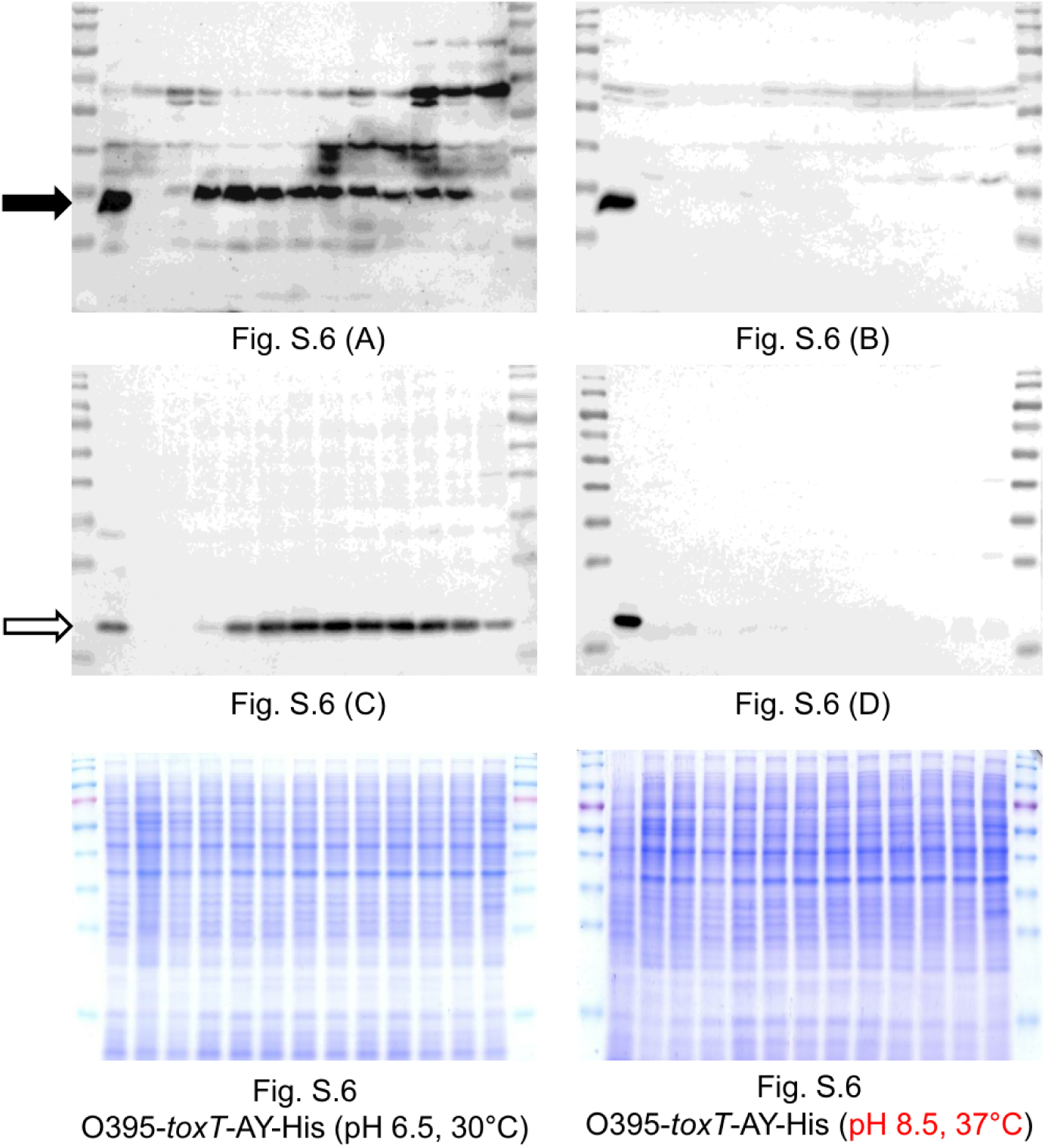
Full size image of western blots and Coomassie brilliant blue-stained SDS-PAGE gels shown in Figures 1–7 and Figures S4–S6.

